# Depletion of the isocitrate dehydrogenase 3 subunit IDHA-1 licenses germ cell-to-neuron direct reprogramming

**DOI:** 10.64898/2026.09.22.753437

**Authors:** Nida ul Fatima, Anna Reid, Amin Shadfar, Alexander Blume, Tobias Opialla, Stefan Kempa, Baris Tursun

**Affiliations:** Department of Biology, Institute of Cell and Systems Biology, Universität Hamburg, 20146 Hamburg, Germany; Max-Delbrück Center for Molecular Medicine, Berlin Institute for Medical Systems Biology, 10115 Berlin, Germany

**Keywords:** direct reprogramming, cell fate, germ line, metabolic reprogramming barrier, *C. elegans*, TCA cycle, isocitrate dehydrogenase, α-Ketoglutarate, HIF-1, anaplerosis

## Abstract

Direct reprogramming (DR) converts one differentiated cell identity into another and is a promising strategy for regenerative approaches when cells are lost because of injury or disease. Transcription factors (TFs) that specify a given cell type can drive DR, but inhibitory mechanisms restrict this conversion in most cell types. To identify such barriers *in vivo*, we use *Caenorhabditis elegans* and the zinc finger TF CHE-1, which is required to specify the glutamatergic taste neuron fate of ASE neurons. Ectopic CHE-1 expression can induce DR of different cell types to ASE neurons upon RNAi-mediated depletion of barrier genes.

Here we characterize the α subunit of the mitochondrial isocitrate dehydrogenase 3 complex (IDH3), IDHA-1, as a barrier to DR of germ cells into neurons. RNAi against *idha-1* produced consistent ASE fate reporter expression, accompanied by morphological changes to neuron-like structures in the germline upon ectopic CHE-1 expression. Using different neuronal gene expression reporters, single-molecule FISH, and antibody staining, we confirm that this germ cell conversion (GeCo) produces neuron-like cells. We found that loss of IDH3 activity causes metabolic perturbations that result in a variety of direct and indirect effects on DR of germ cells. One arm implicates the hypoxia-inducible factor HIF-1, which promotes GeCo. Its loss causes a significant decrease, while the *vhl-1* mutant background, in which HIF-1 protein is stabilized, leads to enhanced GeCo. Another arm implicates epigenetic changes leading to a detectable loss of the repressive histone marks H3K27me3 and H3K9me3 upon *idha-1* depletion. Furthermore, stable-isotope-resolved metabolomics shows strong citrate accumulation without a noteworthy drop in α-Ketoglutarate (αKG) levels, indicating that compensatory pathways maintain αKG levels. Genetics and metabolomics analyses confirmed that Glutamate anaplerosis contributes to compensating for the loss of IDHA-1. Additionally, we found that depletion of glucose transporter FDGT-2 nearly doubles GeCo efficiency. Overall, our findings define the TCA cycle as an *in vivo* safeguard of germ cell identity and show that multiple metabolic inputs converge to keep germ cells refractory to TF-induced DR to neuronal cells.

## INTRODUCTION

Differentiated cell identity is maintained by mechanisms that prevent activation of alternative gene expression programs. Yet, transcription factors (TFs) can revert differentiated cells to pluripotency by reprogramming, or convert them directly into another differentiated type (Davis, Weintraub et al. 1987, Schneuwly, Klemenz et al. 1987, Takahashi and Yamanaka 2006, Vierbuchen, Ostermeier et al. 2010). Cellular reprogramming is typically inefficient or completely blocked, indicating that barriers safeguard resident identity. Hence, to induce cell fate conversion, a TF must overcome the cell’s molecular processes that restrict reprogramming (Soufi and Zaret 2013, Becker, Nicetto et al. 2016, Brumbaugh, Di Stefano et al. 2019). Identifying these barriers helps clarify how cellular identity is maintained and can make direct reprogramming (DR) more efficient for regenerative applications. Previously characterized barriers often relate directly to chromatin regulation and include histone chaperones, chromatin remodellers, histone-modifying enzymes, and complexes that maintain chromatin states. They have repeatedly emerged from unbiased screens for barriers using TF-induced fate conversion in both invertebrate and vertebrate systems (Tursun, Patel et al. 2011, Onder, Kara et al. 2012, Cheloufi, Elling et al. 2015, Kolundzic, Ofenbauer et al. 2018, Hajduskova, Baytek et al. 2019). A less explored class of molecular processes that affects cell identity is cellular metabolism. Reprogramming somatic cells into induced pluripotent stem cells (iPSCs) shifts metabolism from oxidative phosphorylation to glycolysis, and manipulating this shift alters reprogramming efficiency (Folmes, Nelson et al. 2011, Zhang, Nuebel et al. 2012, Zhang, Wang et al. 2015). Metabolism is mechanistically linked to chromatin state by delivering substrates or cofactors of chromatin-modifying enzymes such as acetyl-CoA for histone acetyltransferases, S-adenosylmethionine for methyltransferases, and α-Ketoglutarate (αKG) for dioxygenases that include histone demethylases and protein hydroxylases. In particular, αKG which is produced in the mitochondrial matrix by isocitrate dehydrogenase, were shown to control the pluripotent state of mouse embryonic stem cells (Carey, Finley et al. 2015, Tischler, Gruhn et al. 2019) accelerate differentiation of primed human pluripotent stem cells (TeSlaa, Chaikovsky et al. 2016), and prevent transformation of fibroblasts into cancer-associated fibroblasts (Zhang, Wang et al. 2015). Also, succinate and fumarate act as competitive inhibitors of αKG-dependent dioxygenases thereby blocking cellular transformation to glioma and leukemia (Figueroa, Abdel-Wahab et al. 2010, Xu, Yang et al. 2011, Lu, Ward et al. 2012, Xiao, Yang et al. 2012). Whether perturbations at the αKG node can create permissiveness for direct reprogramming *in vivo* remains unclear because it requires a system in which fate conversion can be assessed quantitatively in an intact animal.

The nematode *C. elegans* serves as such a model system, providing a transparent body for fluorescent fate reporters with efficient RNAi and other molecular techniques. We previously performed a whole-genome RNAi screen for candidate barrier genes using the ectopic expression of the Zn-finger TF CHE-1 (Kolundzic, Ofenbauer et al. 2018). CHE-1 is the terminal selector of the glutamatergic ASE taste neuron (Chang, Johnston et al. 2003, Etchberger, Lorch et al. 2007), but is alone insufficient to convert cells when overexpressed in other tissues. However, upon depletion of the histone chaperone LIN-53 (RBBP4/7) or FACT, germ cells are converted into ASE-like neurons that express the ASE marker *gcy-5* and adopt neuron-like morphology (Tursun, Patel et al. 2011, Kolundzic, Ofenbauer et al. 2018). Removal of Polycomb repressive complex 2 components phenocopies this effect (Patel, Tursun et al. 2012), and subsequent work identified the chromodomain protein MRG-1 (Hajduskova, Baytek et al. 2019) as further germ cell identity safeguards. Importantly, the whole-genome RNAi screen recovered not only chromatin factors but also genes with no obvious connection to chromatin and gene regulation. Among them was *idha-1*, encoding the α subunit of the mitochondrial NAD-dependent isocitrate dehydrogenase 3 complex (IDH3) (Lin, Chang et al. 2022). RNAi against *idha-1* allows overexpressed CHE-1 to induce the ASE fate reporter *gcy-5::gfp* in the germline, raising the possibility that the TCA cycle contributes to germ cell identity maintenance *in vivo*. Here, we show that depletion of the catalytic α or the regulatory γ subunit of IDH3, but not other isocitrate dehydrogenase genes, renders germ cells permissive for CHE-1-induced conversion into neuron-like cells.

Perturbing a central metabolic enzyme such as an isocitrate dehydrogenase can trigger multiple changes beyond altered TCA intermediates and amino acid pools, including changes in oxygen sensing, nuclear histone methylation, and many other cellular processes. Hence, the causal chain from loss of IDH3 activity to the permissive state is not straightforward to establish. Our data show that GeCo upon *idha-1* depletion is promoted by HIF-1 and specific JmjC-domain proteins, and accompanied by loss of repressive histone methylation in the germline. Surprisingly, knockdown of *idha-1* is not accompanied by a measurable decrease in bulk αKG but by the modulation of glutamine/glutamate anaplerosis and glucose transport. We therefore present a model in which several partially independent metabolic inputs converge on germline chromatin to allow TF-induced direct reprogramming.

## RESULTS

### Depletion of the IDH3 α or γ subunit permits CHE-1-induced germ cell conversion

To identify reprogramming barriers *in vivo*, we previously established animals carrying a heat-shock-inducible broad overexpression transgene of *che-1* together with the ASE fate reporter *gcy-5^prom^::GFP* (Tursun, Patel et al. 2011). CHE-1 overexpression alone is very limited in inducing *gcy-5^prom^::GFP* , unless depletion of a barrier gene lifts this restriction (Tursun, Patel et al. 2011, Kolundzic, Ofenbauer et al. 2018) (Figure 1A). The gene *idha-1,* encoding a mitochondrial isocitrate dehydrogenase, was identified in a previous genome-wide RNAi screen (Kolundzic, Ofenbauer et al. 2018) but remained uncharacterized. Combining *idha-1* RNAi with CHE-1 overexpression (CHE-1oe) induced *gcy-5^prom^::GFP* expression in the germline, accompanied by morphological changes in which germ cells acquired neuron-like structures resembling axo-dendritic projections (Figure 1B).

**Figure 1.**
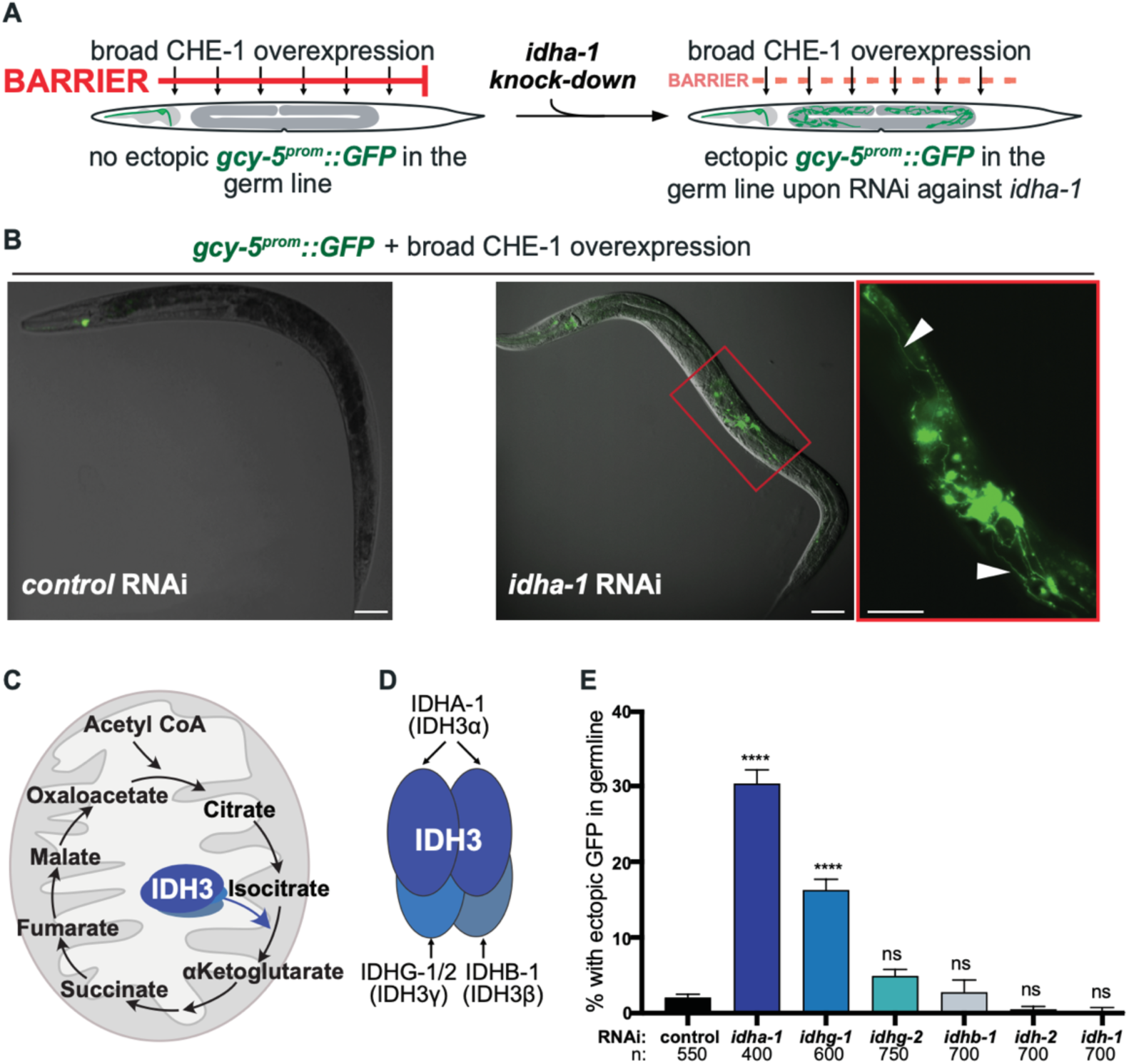
Depletion of mitochondrial IDH3 subunits permits CHE-1-induced ASE fate reporter expression in the *C. elegans* germline. **(A)** Schematic of the transcription factor-induced direct reprogramming system. Animals express the ASE fate reporter *gcy-5^prom^*::*GFP* in the ASE neuron only. Broad overexpression of the ASE neuron fate-inducing transcription factor CHE-1 (CHE-1^OE^) is insufficient to induce the ASE fate reporter in additional cells, indicating cell-fate safeguarding mechanisms that restrict ectopic CHE-1 activity. RNAi-mediated depletion of *idha-1* decreases the barrier and permits ectopic *gcy-5*^prom^::*GFP* expression in germ cells. **(B)** Representative images of animals carrying *gcy-5*^prom^::*GFP* and subjected to broad CHE-1 overexpression following control or *idha-1* RNAi. Ectopic GFP expression is absent from the germline in control RNAi-treated animals but is induced in germ cells following *idha-1* depletion. These *gcy-5*^prom^::*GFP*-positive germ cells also acquire a neuron-like morphology, including prominent neurite-like projections that are clearly distinct from the typical morphology of germ cells. Boxed regions are shown at higher magnification; arrowheads indicate ectopic *gcy-5*^prom^::*GFP* positive germ cells and associated projections. **(C)** Schematic representation of the mitochondrial tricarboxylic acid (TCA) cycle. NAD+-dependent isocitrate dehydrogenase 3 (IDH3) catalyzes the conversion of isocitrate to α-ketoglutarate (α-KG). **(D)** Schematic of the heterotetrameric mammalian IDH3 complex, comprising two α-subunits, one β-subunit and one γ-subunit, with the corresponding *C. elegans* homologs IDHA-1, IDHB-1 and IDHG-1 indicated. **(E)** Quantification of animals displaying ectopic *gcy-5*^prom^::*GFP* expression in the germline following RNAi-mediated depletion of the indicated IDH3 subunits and related isocitrate dehydrogenases in combination with CHE-1^OE^. Depletion of *idha-1* or *idhg-1*, encoding the α and γ subunits of mitochondrial NAD^+^-dependent IDH3, respectively, significantly increased ectopic germline GFP expression relative to control RNAi. In contrast, depletion of the isocitrate dehydrogenases *idh-1* and *idh-2* did not permit ectopic germline GFP expression. IDH-1 and IDH-2 are NADP+-dependent isocitrate dehydrogenases, distinct from the NAD+-dependent IDH3 complex that catalyses the canonical oxidative TCA cycle reaction. Data are presented as mean ± SEM from three independent biological experiments. Statistical significance was determined by ordinary one way ANOVA followed by Dunnett’s multiple comparisons test, with each RNAi condition compared with control RNAi. ****P < 0.0001; ns, not significant. Numbers below the bars indicate the total number of animals scored.

IDHA-1 protein is part of the NAD⁺-dependent isocitrate dehydrogenase complex IDH3 of the mitochondrial matrix, which catalyzes the oxidative decarboxylation of isocitrate to αKG, in the TCA cycle (Gabriel, Zervos et al. 1986) (Figure 1C). In humans, the enzyme assembles as a heterotetramer of catalytic α subunits with regulatory β and γ subunits (Chen and Ding 2023); *C. elegans* genes encode one α subunit (IDHA-1), one β subunit (IDHB-1), and two γ subunits (IDHG-1 and IDHG-2) (Figure 1D). We tested RNAi against the corresponding genes and those encoding other isocitrate dehydrogenase genes, including the NADP-dependent enzymes IDH-1 and IDH-2. Only depletion of *idha-1* (∼30% of animals; n = 400) and *idhg-1* (∼16%; n = 600) allowed *gcy-5^prom^::GFP* induction by CHE-1oe (Figure 1E). The barrier effect is therefore specific to the α and one of the two γ subunits of the mitochondrial NAD-dependent complex.

To assess which tissues IDHA-1 and IDHG-1 are expressed, we generated CRISPR knock-in alleles tagging IDHA-1 with 3xHA and IDHG-1 with 3xFLAG for immunostaining. Both proteins were broadly expressed across tissues, including the germline, showing an intracellular pattern characteristic of mitochondria based on MitoTracker staining (Suppl.Figure 1). However, we could not generate homozygous *idha-1* null mutants using CRISPR/Cas9-mediated genomic deletion. While heterozygous animals showed mild developmental defects, their progeny homozygous for the *idha-1* deletion mutation died as embryos or early larvae. Therefore, we conducted all subsequent experiments using RNAi to knock down *idha-1*.

### Converted germ cells express additional neuronal genes

Because ectopic expression of a single reporter does not reflect a faithful change in cell identity, we tested whether converted germ cells activate neuronal genes beyond *gcy-5*, at both subtype-specific and pan-neuronal levels. For clarity, we refer to the CHE-1-induced germ cell conversion hereafter as GeCo (<u>Ge</u>rm <u>Co</u>nversion).

We combined *gcy-5^prom^::GFP* with RFP reporters for three additional identity levels: *ceh-36^prom^::RFP* (the ASE/AWC-class homeobox gene acting downstream of *che-1* in ASE specification), *ift-20^prom^::NLS::RFP* (a sensory/ciliated neuron marker), and *rab-3^prom^::NLS::RFP* (pan-neuronal). All three reporters were induced in converted germ lines of *idha-1* RNAi animals (Figure 2A). Quantification showed that around 60% of animals with germline GFP also expressed either *ceh-36^prom^::RFP*, *rab-3^prom^::RFP*, or *ift-20^prom^::RFP* (Figure 2B). Hence, CHE-1 induced GeCo co-activates ASE-specific, sensory, and pan-neuronal reporter gene expression, arguing that a broader neuronal program rather than a single promoter is being induced.

**Figure 2.**
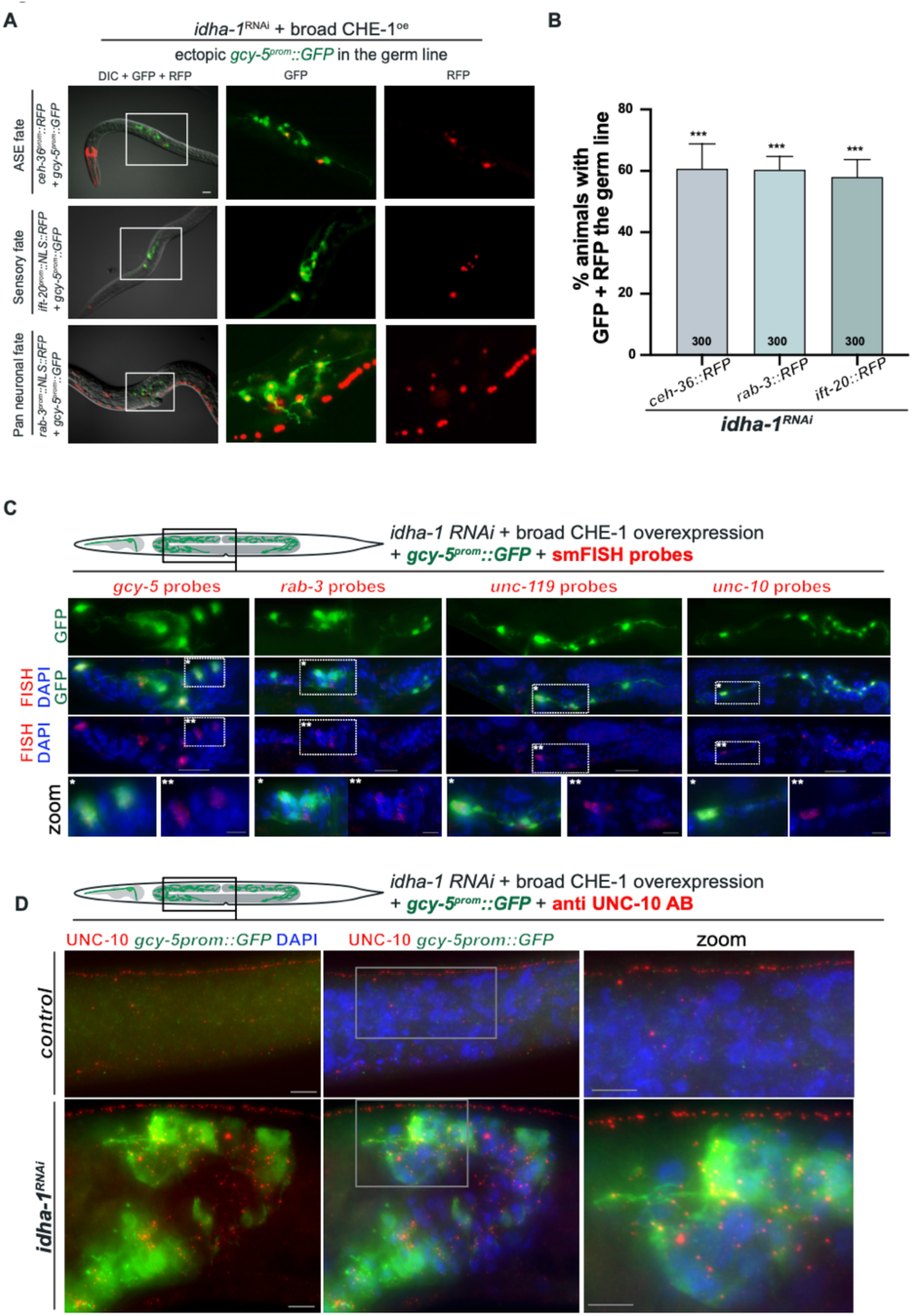
CHE-1-reprogrammed germ cells acquire neuronal characteristics. **(A)** Representative images of ectopic neuronal reporter expression in the germline following *idha-1* RNAi and broad CHE-1^OE^. Animals expressed the ASE fate reporter *gcy-5*^prom^::*GFP* with reporters for additional neuronal identity markers: *ceh-36*^prom^::*RFP*, an ASE/AWC neuronal fate marker; *ift-20*^prom^::*NLS::RFP* (marker of ciliated sensory neurons); and the pan-neuronal reporter *rab-3*^prom^::*NLS::RFP*. For each reporter combination, merged DIC/GFP/RFP, GFP/RFP, and RFP channels are shown. Boxed regions indicate areas containing ectopic reporter-positive germ cells. These additional neuronal reporters were detected within *gcy-5*^prom^::*GFP* positive germ cells following *idha-1* depletion and CHE-1^OE^. **(B)** Quantification of animals displaying co-expression of ectopic *gcy-5*^prom^::*GFP* and the indicated neuronal RFP reporter in the germline following *idha-1* RNAi and CHE-1^OE^. Approximately 60% of animals showed co-expression of each neuronal reporter with *gcy-5*^prom^::*GFP*, compared with ∼3% in controls. **Data are presented as mean ± SEM. Statistical significance was determined by ordinary one-way ANOVA followed by Dunnett’s multiple comparisons test, with each condition compared with control. ***P < 0.001. Numbers within the bars indicate the total number of animals scored. **(C)** Detection of endogenous neuronal transcripts in reprogrammed germ cells by single-molecule fluorescence *in situ* hybridization (smFISH). Animals subjected to *idha-1* RNAi and CHE-1^OE^ and displaying ectopic *gcy-5* ^prom^::*GFP* expression were hybridized with probes targeting endogenous *gcy-5, rab-3, unc-119,* or *unc-10*. GFP marks *gcy-5* ^prom^::*GFP*-positive germ cells; smFISH signals are shown as red puncta, and nuclei are counterstained with DAPI (blue). Boxed regions are shown at higher magnification in the bottom row. Endogenous transcripts for all four neuronal genes were detected within ectopic GFP-positive germ cells. **(D)** Immunofluorescence detection of the conserved presynaptic protein UNC-10/RIM in the germline. Representative control and *idha-1* RNAi-treated animals following CHE-1^OE^ are shown. UNC-10 immunostaining is shown in red, *gcy-5* ^prom^::*GFP* in green, and DAPI in blue. Ectopic UNC-10 protein is detected within *gcy-5* ^prom^::*GFP*-positive germ cells following *idha-1* depletion, whereas comparable neuronal marker expression is absent from the control germline. Boxed regions are shown at higher magnification. **Short conclusion:** Together, these results show that *idha-1* depletion permits CHE-1 to activate a broader neuronal program in germ cells and supports the acquisition of a broader neuronal molecular identity rather than isolated activation of the *gcy-5* fate reporter.

We confirmed that the transgenic reporters reflect genuine gene expression by examining endogenous transcripts using single-molecule fluorescence *in situ* hybridization (smFISH). In GFP-positive germlines of *idha-1* RNAi animals, we detected smFISH signals for *gcy-5* as well as the pan-neuronal genes *rab-3*, *unc-119,* and *unc-10* (Figure 2C). Furthermore, antibody staining against the neuronal active-zone protein UNC-10/RIM revealed punctate UNC-10 signal within *gcy-5^prom^::GFP*-positive germ cells of *idha-1* RNAi animals, whereas control animals showed only the expected UNC-10 staining in the neighboring nerve cord and none in the germ line (Figure 2D). Together, these data show that IDHA-1 depletion and ectopic CHE-1oe activate endogenous neuronal genes at the transcript and protein level in germ cells that acquire neuron-like morphology, corroborating the notion of genuine germ cell-to-neuron conversion.

We also asked whether the permissive state created by *idha-1* depletion is specific to the ASE neuron fate. Ectopic expression of the Pitx-type homeodomain TF UNC-30, which specifies GABAergic motor neurons (Jin, Hoskins et al. 1994), induced the GABA transporter reporter *unc-25^prom^::GFP* in the germline of *idha-1* RNAi animals. However, ectopic expression of the myogenic bHLH factor HLH-1 induced no germline muscle reporter expression (Suppl.Figure 2). Hence, loss of IDHA-1 lowers a barrier that restricts conversion towards at least two distinct neuronal identities but does not seem to erase germ cell identity per se to allow induction of the muscle fate by HLH-1.

### Loss of IDHA-1 reduces repressive histone methylation in the germline

Because αKG is the co-substrate of JmjC-domain histone demethylases, we asked whether depletion of IDHA-1 alters the histone methylation landscape. Western blotting of whole-worm lysates from *idha-1* RNAi-treated animals for histone marks revealed essentially unchanged H3K4me3, which marks active chromatin and a modest reduction of the repressive histone marks H3K27me3 and H3K9me3 (Figure 3A). To validate IDHA-1 knockdown, we used the endogenous IDHA-1::3xHA allele for Western blotting of whole-worm lysates, with tubulin and 14-3-3 as loading controls (Figure 3A). Since whole-animal lysates contain mixed tissue including somatic and germ cells, we examined the germline directly by immunofluorescence, using HTZ-1/H2A.Z counterstaining for normalization (Figure 3B). Relative to control RNAi, *idha-1* RNAi reduced germline H3K27me3 more pronouncedly, while H3K4me3 was slightly increased (Figure 3C). Thus, *idha-1* knockdown reduces repressive histone marks, the opposite of what αKG limitation should cause if reduced αKG reduces JmjC activity that demethylates H3K9/27me3. Yet the observed loss of H3K9me3 and H3K27me3 upon *idha-1* depletion explains the increased permissiveness for ectopic gene expression in germlines, as we observed in previous studies of barrier genes (Tursun, Patel et al. 2011, Patel, Tursun et al. 2012).

**Figure 3.**
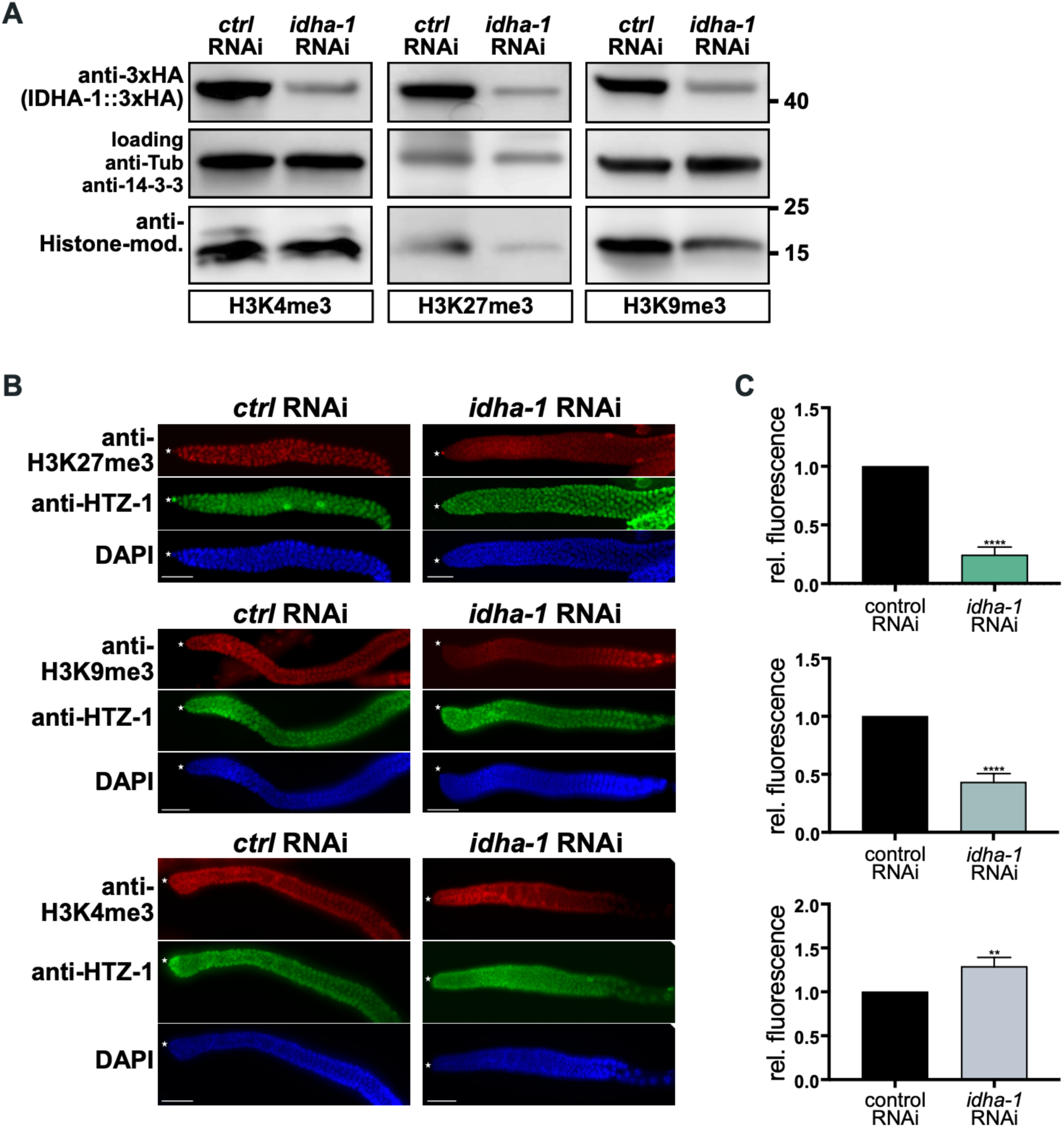
IDHA-1 depletion remodels histone methylation in the *C. elegans* germline. **(A)** Whole-worm western blot analysis of histone methylation following RNAi-mediated depletion of *idha-1*. IDHA-1 depletion was confirmed in animals carrying endogenously tagged IDHA-1::3xHA using anti-HA antibodies. Whole-animal protein lysates from control and *idha-1* RNAi-treated animals were probed for the active chromatin-associated histone modification H3K4me3 and the repressive histone modifications H3K27me3 and H3K9me3. α-tubulin/14-3-3 was used as loading controls. Depletion of *idha-1* reduced H3K27me3 and H3K9me3 levels, indicating loss of repressive histone methylation following perturbation of mitochondrial IDH3. **(B)** Representative immunofluorescence images of germ lines from control and *idha-1* RNAi-treated animals stained for H3K27me3, H3K9me3, or H3K4me3 (red). HTZ-1/H2A.Z staining (green) was used as a chromatin reference, and nuclei were counterstained with DAPI (blue). *idha-1* depletion markedly reduced germline H3K27me3 and H3K9me3 fluorescence. Asterisks indicate the distal tip of the germline. Scale bars, 50 μm. **(C)** Quantification of relative fluorescence intensities of H3K27me3, H3K9me3, and H3K4me3 in germ lines from control and *idha-1* RNAi-treated animals. *idha-1* depletion markedly reduced the repressive histone modifications H3K27me3 and H3K9me3, accompanied by a significant increase in the active chromatin-associated mark H3K4me3. Data are presented as mean ± SEM. ****P < 0.0001; **P < 0.01.

This outcome suggests that chromatin state is not directly affected by reduced demethylase co-substrate availability and requires the genetic and metabolomic dissection of effects caused by *idha-1* depletion described below.

### GeCo requires HIF-1 and the JmjC-domain proteins JMJD-3.3 and JMJD-4

Knockdown of *idha-1* is expected to interrupt the TCA cycle at the isocitrate-to-αKG step, which could create permissiveness for CHE-1-driven GeCo through several routes (Figure 4A). Besides JmjC oxygenase-mediate histone demthylation, another prominent αKG-dependent pathway is the hypoxia response: the prolyl hydroxylase EGL-9 (PHD) requires αKG to hydroxylate HIF-1 which then serves as signal for VHL-1-dependent ubiquitylation and proteasomal degradation (Figure 4B) (Epstein, Gleadle et al. 2001, Kaelin and Ratcliffe 2008).

**Figure 4.**
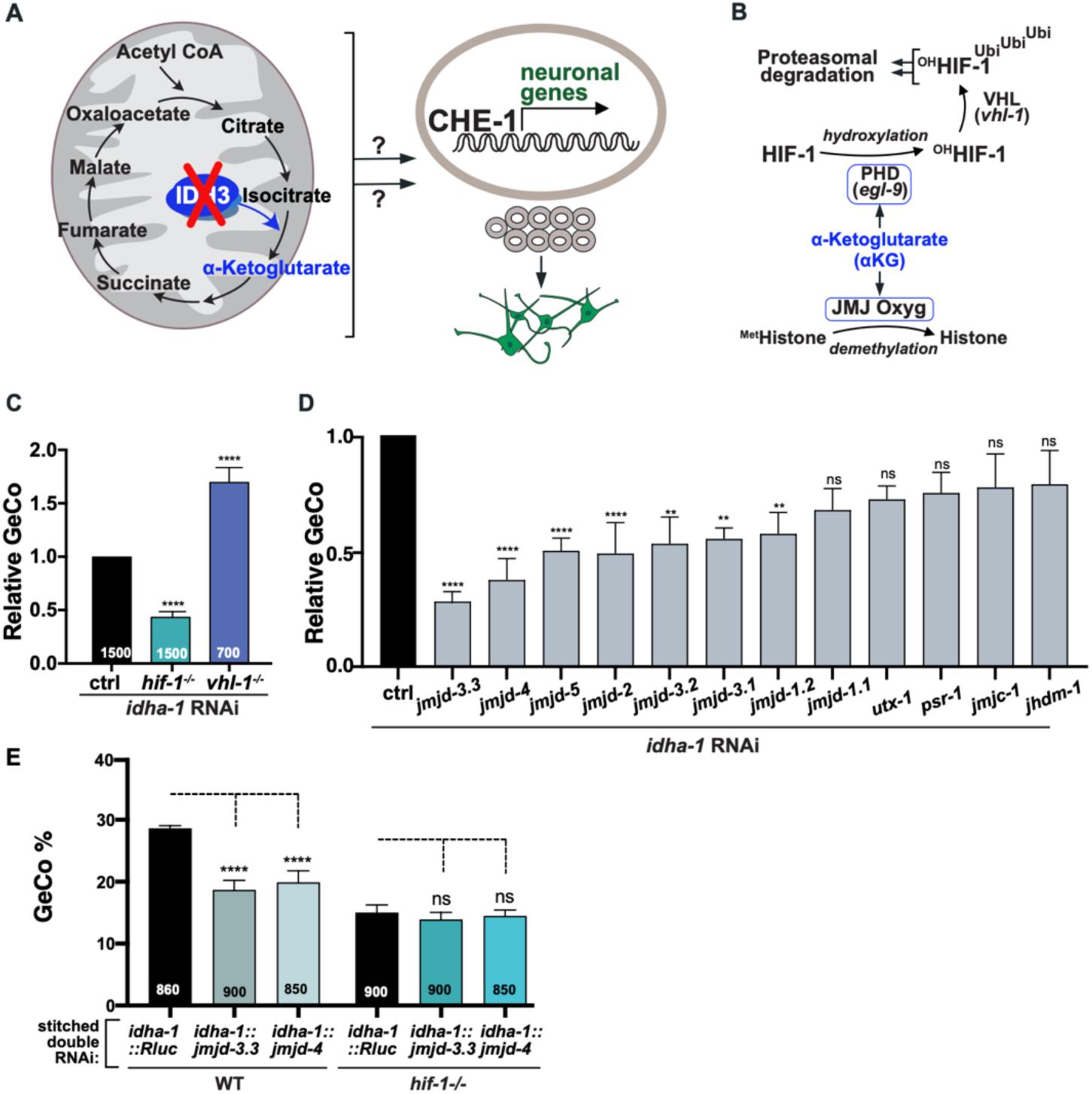
HIF-1 and Jumonji histone demethylases promote IDHA-1 depletion-mediated germ cell reprogramming. **(A)** Working model for mitochondrial regulation of transcription factor-induced germ cell reprogramming. Depletion of the mitochondrial TCA-cycle enzyme IDH3 through *idha-1* RNAi is proposed to trigger retrograde effects from mitochondria to the nucleus, altering the chromatin environment and enabling ectopic CHE-1 to activate a neuronal gene program in germ cells. Question marks indicate unknown links between mitochondrial IDH3 depletion and nuclear changes. **(B)** Schematic of α-KG-dependent pathways linking IDH3 function to HIF-1 stability and chromatin regulation. The prolyl hydroxylase EGL-9/PHD uses α-KG for HIF-1 hydroxylation, enabling VHL-1-dependent ubiquitination and proteasomal degradation of HIF-1. α-KG is also a cofactor for Jumonji-domain oxygenases that catalyse histone demethylation. **(C)** Genetic modulation of HIF-1 signaling alters IDHA-1 depletion-mediated germ cell conversion (GeCo). Relative GeCo penetrance following *idha-1* RNAi was quantified in control, *hif-1*^−/−^, and *vhl-1*^−/−^ animals following CHE-1^OE^. Loss of *hif-1* strongly suppressed the *idha-1*-dependent reprogramming phenotype, whereas loss of *vhl-1* enhanced GeCo, consistent with HIF-1 activity promoting reprogramming following IDH3 depletion. Statistical significance was determined by ordinary one-way ANOVA followed by Dunnett’s multiple-comparisons test, with mutant conditions compared with control. ****P < 0.0001. **(D)** RNAi suppressor screen for Jumonji-domain proteins required for IDHA-1 depletion-mediated reprogramming. Candidate Jumonji genes were co-depleted with *idha-1*, and GeCo penetrance following CHE-1^OE^ was normalized to the *idha-1* RNAi control. Co-depletion of *jmjd-3.3, jmjd-4, jmjd-5, jmjd-2, jmjd-3.2, jmjd-3.1,* and *jmjd-1.2* significantly suppressed the reprogramming phenotype, whereas depletion of *jmjd-1.1, utx-1, psr-1, jmjc-1,* or *jhdm-1* did not significantly alter GeCo. The candidates *jmjd-3.3* and *jmjd-4* were selected for further validation. Statistical significance was determined by ordinary one-way ANOVA followed by Dunnett’s multiple-comparisons test, with each double-RNAi condition compared with the *idha-1* RNAi control. ****P < 0.0001; **P < 0.01; ns, not significant. **(E)** Genetic interaction between HIF-1 and the selected candidate Jumonji factors *jmjd-3.3* and *jmjd-4*. Stitched double-RNAi constructs were used to simultaneously deplete *idha-1* and either *jmjd-3.3* or *jmjd-4*, with *idha-1::Rluc* RNAi used as the corresponding control. In wild-type animals, co-depletion of either *jmjd-3.3* or *jmjd-4* significantly reduced GeCo compared with *idha-1::Rluc* RNAi. In the *hif-1*^−/−^ background, however, additional depletion of *jmjd-3.3* or *jmjd-4* did not further suppress the reprogramming phenotype, consistent with HIF-1 and these Jumonji factors acting within the same genetic pathway. Data are presented as mean ± SEM. Numbers within bars indicate the total number of animals scored. Statistical significance was determined by ordinary one way ANOVA followed by Tukey’s multiple comparisons test. ****P < 0.0001; ns, not significant.

Using a viable *hif-1* null mutant background, we observed that GeCo upon *idha-1* RNAi was significantly reduced by approximately 50%. Conversely, *vhl-1* null animals, in which HIF-1 protein is constitutively stabilized (Bishop, Lau et al. 2004, Shen, Nettleton et al. 2005), showed strong GeCo enhancement by 1.5-fold (Figure 4C). HIF-1 activity appears to be rate-limiting but also to promote GeCo upon *idha-1* depletion. This effect is not caused by altered levels of the reprogramming-driving TF CHE-1, as Western blotting of heat-shocked animals carrying CHE-1::3xHA showed equal levels of induced CHE-1 protein in wild-type and *hif-1* mutant backgrounds (Suppl.Figure 3A).

The observation of reduced histone methylation (Figure 3B,C) suggested that αKG-dependent dioxygenases that mediate histone demethylation may be required for CHE-1-induced GeCo. We screened the *C. elegans* JmjC-domain gene family by double RNAi combined with *idha-1* knockdown and found that co-depletion of *jmjd-3.3* and *jmjd-4* most strongly suppressed GeCo, while a second tier of genes, including *jmjd-5*, *jmjd-2*, *jmjd-3.2*, *jmjd-3.1,* and *jmjd-1.2*, gave statistically significant but weaker suppression (Figure 4F). The available mutant alleles for *jmjd-3.3* and *jmjd-4* showed weaker suppression than RNAi (Suppl. Figure 3B), which we attribute to genetic compensation within this partially redundant family; the two available *jmjd-3.3* alleles are short in-frame deletions that may preserve at least partial protein activity. We therefore followed up on *jmjd-3.3* and *jmjd-4* with stitched double-RNAi clones, which can be used for more stable double knockdown assessments (Kazmierczak, Farre et al. 2021). The *idha-1* RNAi clone fragment is fused in the L4440 RNAi vector with a *jmjd-3.3* or *jmjd-4* genomic fragment, or with the *Rluc* gene fragment as the matched control. Stitched *idha-1::jmjd-3.3* reduced GeCo by around 25%, confirming the single-clone result (Figure 4G). Notably, when the same stitched double RNAi was performed in the *hif-1* null background, GeCo was reduced by *idha-1::Rluc* compared to the wild-type background but not further suppressed by co-depletion of *jmjd-3.3* or *jmjd-4* relative to *idha-1::Rluc* in the *hif-1* null background (Figure 4G). Hence, the absence of an additive effect places HIF-1 and these two JmjC proteins in a common genetic pathway for GeCo rather than in parallel pathways. Overall, these findings identify HIF-1 and the JMJDs as central players in creating permissiveness for DR of germ cells in the *idha-1* depletion context. Yet, the directionality of *jmjd-3.3* and *jmjd-4* depletion and their effect on chromatin is not concordant, as reduced demethylase activity should increase histone methylation. This suggests additional or alternative pathways affected by IDH3 activity. Also, because GeCo is not fully abolished upon loss of HIF-1 or JMJD proteins, further effects may be implicated in licensing DR of germ cells to neurons upon *idha-1* depletion.

### Metabolic consequences of *idha-1* depletion

If the permissive state resulted from reduced αKG production, supplying αKG to the *idha-1-depleted* worms should suppress GeCo. We therefore fed animals defined TCA cycle intermediates combined with *idha-1* RNAi and scored GeCo (Figure 5A). Unexpectedly, αKG feeding produced the opposite effect, causing a small enhancement, while malate and fumarate enhanced GeCo by up to 1.5-fold (Figure 5B). Succinate and citrate feeding had no significant effect at the tested concentrations of 10 and 15 mM in the growth medium (Figure 5B). Thus, exogenous supply of the TCA intermediates αKG, malate, and fumarate modulates conversion, but not in the direction that αKG-scarcity would predict.

**Figure 5.**
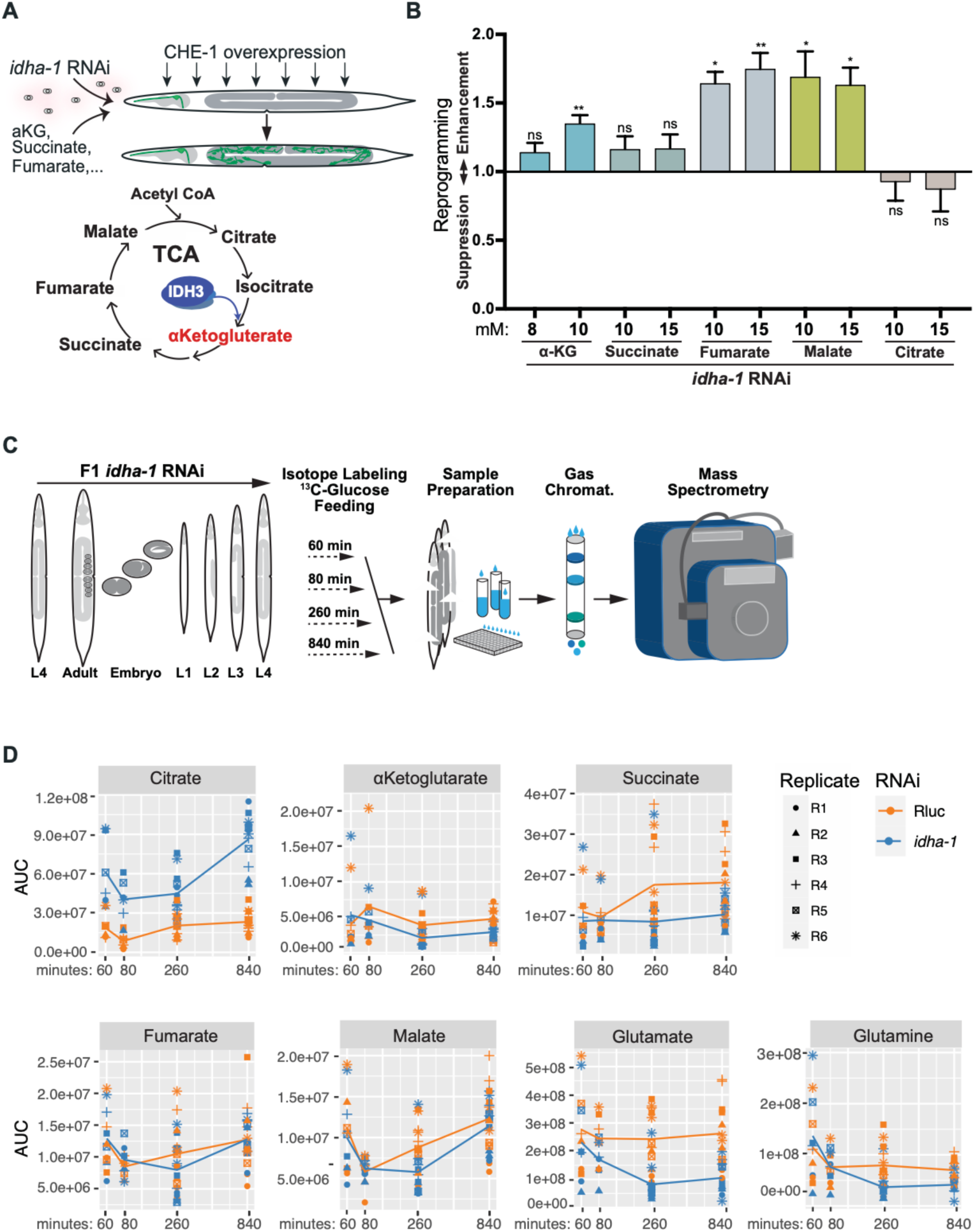
IDHA-1 depletion induces metabolic rewiring. **(A)** Schematic of the metabolite-feeding assay. Animals were treated with *idha-1* RNAi and supplemented with the indicated TCA-cycle metabolites before broad CHE-1^OE^. Germ cell conversion was assessed by ectopic activation of the ASE fate reporter *gcy-5*^prom^::*GFP*. **(B)** Effect of exogenous TCA-cycle metabolites on IDHA-1 depletion-mediated germ cell reprogramming. Animals treated with *idha-1* RNAi were supplemented with α-ketoglutarate (α-KG), succinate, fumarate, malate or citrate at the indicated concentrations, and GeCo penetrance following CHE-1^OE^ was quantified relative to the vehicle-treated *idha-1* RNAi control. α-KG modestly enhanced reprogramming, whereas fumarate and malate produced a stronger increase in GeCo. In contrast, succinate and citrate did not significantly alter the phenotype. Data are presented as mean ± SEM. Statistical significance was determined by one-way ANOVA followed by Dunnett’s multiple comparisons test. **P < 0.01; *P < 0.05; ns, not significant. **(C)** Schematic of the stable isotope tracing experiment used to determine whether carbon flow through the TCA cycle is maintained following IDH3 depletion. F1 animals were exposed to control or *idha-1* RNAi throughout development and subsequently fed ^13^C-labelled glucose. Animals were collected after 60, 180, 360, and 840 min of isotope labeling, followed by metabolite extraction and gas chromatography-mass spectrometry (GC-MS). Time-dependent incorporation of ^13^C into the TCA cycle and associated metabolites was used to assess relative changes in carbon utilization and metabolic pathway activity following *idha-1* depletion. **(D)** Time-resolved ^13^C glucose tracing of TCA-cycle and glutamine-associated metabolites following control (*Rluc*) or *idha-1* RNAi. Despite depletion of the TCA-cycle enzyme IDHA-1, ^13^C incorporation into α-KG, fumarate, and malate remained broadly comparable with control, indicating that downstream TCA-cycle metabolism is at least partially maintained. Citrate showed increased ^13^C labeling following *idha-1* depletion, consistent with accumulation of carbon upstream of the disrupted IDH3-dependent isocitrate-to-α-KG step. Succinate and glutamine showed modest decreases, while glutamate showed a more pronounced reduction in ^13^C labeling following IDH3 depletion, indicating increased utilization of glutamate-associated carbon. Individual symbols represent the 6 independent biological replicates.

We therefore performed metabolomics to directly measure the metabolic consequences of *idha-1* depletion. Using stable-isotope-resolved metabolomics, we exposed P0 L4 animals to *idha-1* or control (*Rluc*) RNAi and maintained their F1 progeny on RNAi until the L4 stage, when animals were fed ¹³C-glucose for 60, 80, 260, or 840 minutes before extraction and GC-MS analysis, with six biological replicates per condition and time point (Figure 5C). The dominant effect was a strong accumulation of citrate in *idha-1*-depleted animals, apparent at 60 minutes and increasing over the time course (Figure 5D), as expected when blocking the isocitrate dehydrogenase step. Unexpectedly, changes downstream of the block were modest, with αKG showing no significant difference between *idha-1* and control RNAi at any time point. Only succinate was modestly reduced at 260 and 840 minutes. However, glutamate was consistently reduced in *idha-1* RNAi (Figure 5D). An independent metabolomics experiment without isotope labeling, comparing P0 and F1 *idha-1* RNAi animals at a single L4 time point, reproduced this pattern, including citrate accumulation and the glutamate decrease (Suppl. Figure 4). These results suggest that GeCo upon *idha-1* depletion occurs without reducing the whole-animal steady-state αKG pool. Yet, this does not exclude a local or compartment-specific change, because GC-MS of whole-worm extracts averages across tissues, and therefore we also attempted GC-MS of dissected gonads from *idha-1* RNAi animals. However, using around 100 dissected germlines as one sample did not provide enough material for even one proper GC-MS run.

Nevertheless, our GC-MS analysis rules out bulk αKG scarcity and is consistent with the observation that αKG feeding fails to suppress conversion. Additionally, the reduction in glutamate indicated altered flux through the glutamine/glutamate node, which prompted us to test this branch genetically.

### Glutamate anaplerosis modulates germ cell conversion

Glutamine and glutamate pools can replenish the TCA cycle at the αKG node by glutaminases (GLNA-1, GLNA-3) hydrolyzing glutamine to glutamate, and by glutamate dehydrogenase (GDH-1) converting glutamate to αKG (Owen, Kalhan et al. 2002, Yang, Ko et al. 2014). Conversely, glutamine synthetase GLN-2 runs the opposite reaction, and αKG can additionally be generated from α-hydroxyglutarate by the hydroxyglutarate dehydrogenases DHGD-1/LHGD-1 (Figure 6A). Testing these possible pathways by double RNAi in the *idha-1* RNAi background showed that glutamine synthetase knockdown by targeting *gln-2*, which is predicted to increase the glutamate pool available for anaplerosis, enhanced GeCo nearly 2-fold (Figure 6B). In contrast, knockdown of the glutaminases GLNA-1 and GLNA-3, as well as of glutamate dehydrogenase GDH-1, strongly suppressed DR of germ cells into neurons. We generated a *glna-3* null mutant which confirms the RNAi phenotype. (Supplemental Figure 5). RNAi against *dhgd-1* or *lhgd-1* had no significant effect (Figure 6B), indicating that the anaplerotic flux from glutamine through glutamate into αKG is required for GeCo. Hence, shifting the balance of this anaplerosis node towards glutamate by removing glutamine synthetase GLN-2 makes germ cells significantly more permissive.

**Figure 6.**
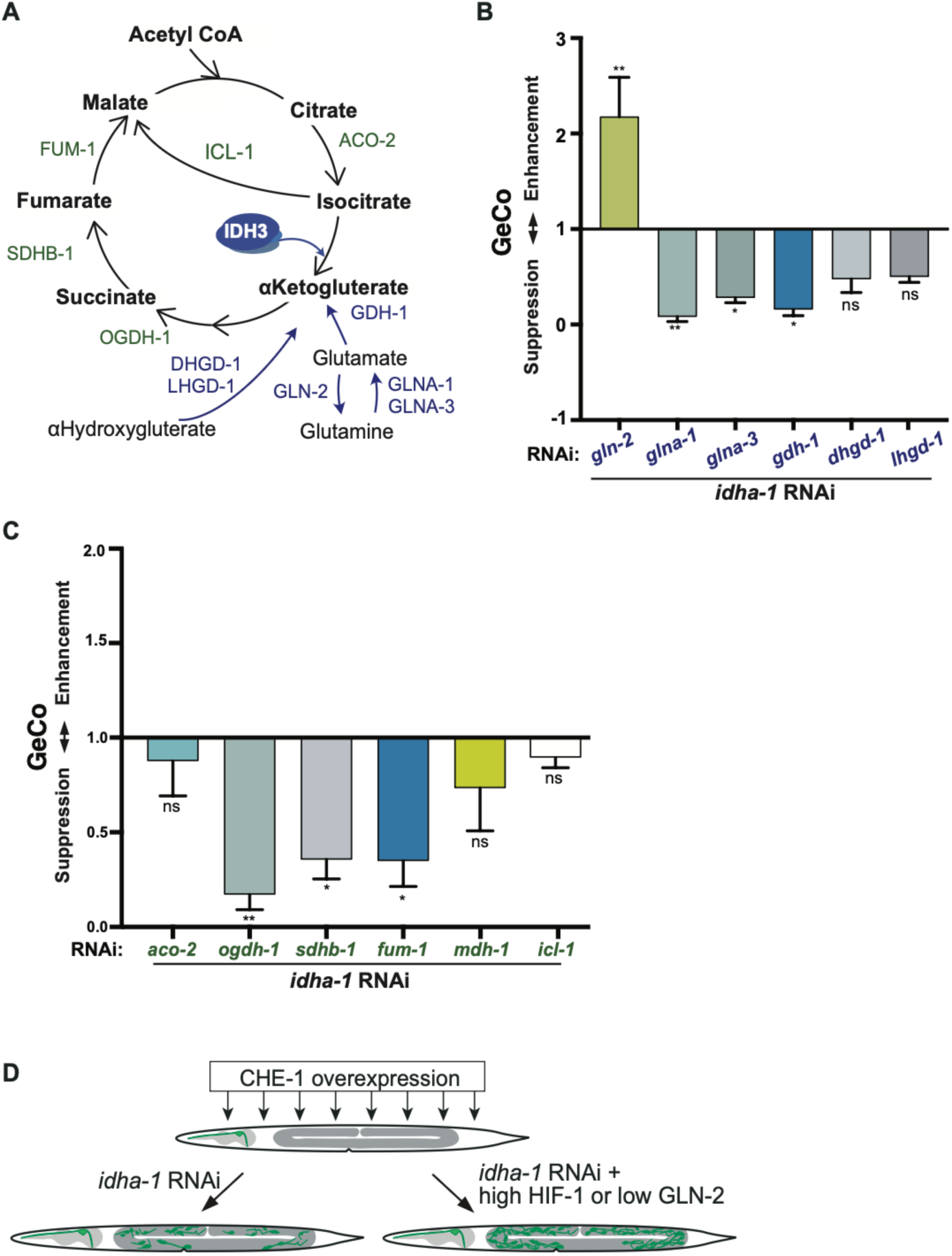
Glutamine anaplerosis and downstream TCA cycle activity support IDH3 depletion-mediated germ cell reprogramming. **(A)** Schematic of metabolic pathways capable of maintaining downstream TCA cycle metabolism following IDH3 depletion. IDH3 normally catalyzes the conversion of isocitrate to α-KG. Alternative α-KG-generating routes include conversion of glutamine to glutamate by the glutaminases GLNA-1 and GLNA-3, followed by conversion of glutamate to α-KG by GDH-1, as well as hydroxyglutarate metabolism. GLN-2 catalyzes the opposing conversion of glutamate to glutamine. The ICL-1-dependent glyoxylate shunt represents an alternative route that could, in principle, maintain TCA cycle function by bypassing the IDH3-dependent isocitrate to α-KG step. Enzymes from these pathways and the downstream TCA cycle were therefore examined genetically. **(B)** Genetic analysis of alternative α-KG generating pathways during IDHA-1 depletion-mediated reprogramming. Candidate metabolic enzymes were co-depleted with *idha-1*, and GeCo penetrance following CHE-1^OE^ was quantified relative to the *idha-1* RNAi control. Depletion of the glutaminases *glna-1* and *glna-3*, or glutamate dehydrogenase *gdh-1*, suppressed reprogramming, whereas depletion of glutamine synthetase *gln-2*, which directs glutamate back toward glutamine, enhanced GeCo. In contrast, depletion of the hydroxyglutarate dehydrogenases *dhgd-1* and *lhgd-1* did not significantly alter the phenotype. These genetic interactions support a functional contribution of glutamate-to-α-KG metabolism during IDH3 depletion. **(C)** Requirement for downstream TCA cycle metabolism during IDHA-1 depletion-mediated reprogramming. TCA cycle and glyoxylate shunt enzymes were co-depleted with *idha-1*, and GeCo was quantified relative to the *idha-1* RNAi control. Co-depletion of *ogdh-1, sdhb-1,* or *fum-1*, which function downstream of α-KG, significantly suppressed reprogramming, whereas depletion of *aco-2, mdh-1*, or the glyoxylate-shunt enzyme *icl-1* did not significantly alter the phenotype. These results indicate that metabolism downstream of α-KG is required for efficient IDH3 depletion-mediated reprogramming, while the ICL-1-dependent glyoxylate shunt is dispensable. **(D)** Working model integrating metabolic and HIF-1-dependent regulation of IDHA-1 depletion-mediated germ cell reprogramming. IDH3 depletion establishes a reprogramming-permissive state that enables CHE-1^OE^ to induce neuronal conversion.**Data are presented as mean ± SEM. Statistical significance was determined by ordinary one-way ANOVA followed by Dunnett’s multiple-comparisons test, with each double-RNAi condition compared with the *idha-1* RNAi control. ***P < 0.001; **P < 0.01; *P < 0.05; ns, not significant.

Consistent with a requirement for glutamate handling rather than for glutamine itself, glutamine feeding had no significant effect on GeCo, whereas RNAi against the glutamate transporter GLFT-1 reduced conversion (Supplemental Figure 5).

We also tested whether depletion of TCA enzymes acting downstream of the *idha-1* block affected GeCo. Depletion of the αKG dehydrogenase OGDH-1, the succinate dehydrogenase subunit SDHB-1, and the fumarase FUM-1 each strongly suppressed GeCo, whereas depletion of the aconitase ACO-2, the malate dehydrogenase MDH-1, and the isocitrate lyase ICL-1 had no significant effect (Figure 6D). Thus, GeCo requires continued flux from αKG to malate and is triggered by loss of flux at the isocitrate-to-αKG step. This is consistent with the metabolite-feeding data, in which fumarate and malate enhanced rather than suppressed conversion.

Taken together, these results reveal two conditions caused by *idha-1* depletion that contribute to the permissiveness of germ cells DR to neurons: elevated HIF-1 and glutamate anaplerosis levels (Figure 6E).

### The putative glucose transporter FDGT-2 is an additional GeCo barrier

To systematically search for additional metabolic modifiers, we performed an extended double RNAi screen with *idha-1* RNAi combined with ∼110 genes encoding metabolic enzymes, solute carriers, and transporters, including the glutamate anaplerosis and TCA genes tested above as internal controls (Figure 7A). The screen reproduced these and also identified a new target: RNAi against *fdgt-2*, which encodes a facilitated glucose transporter of the GLUT/SLC2A family (Kim, Underwood et al. 2018), markedly increased conversion relative to *idha-1* RNAi alone (Figure 7A). We confirmed this finding with a CRISPR-generated *fdgt-2* null mutant, in which GeCo upon *idha-1* RNAi was enhanced almost 2-fold over *idha-1* depletion alone (Figure 7B). We also generated an FDGT-2::3xHA knock-in allele to assess expression and could detect broad expression across tissues (Supplemental Figure 6).

**Figure 7.**
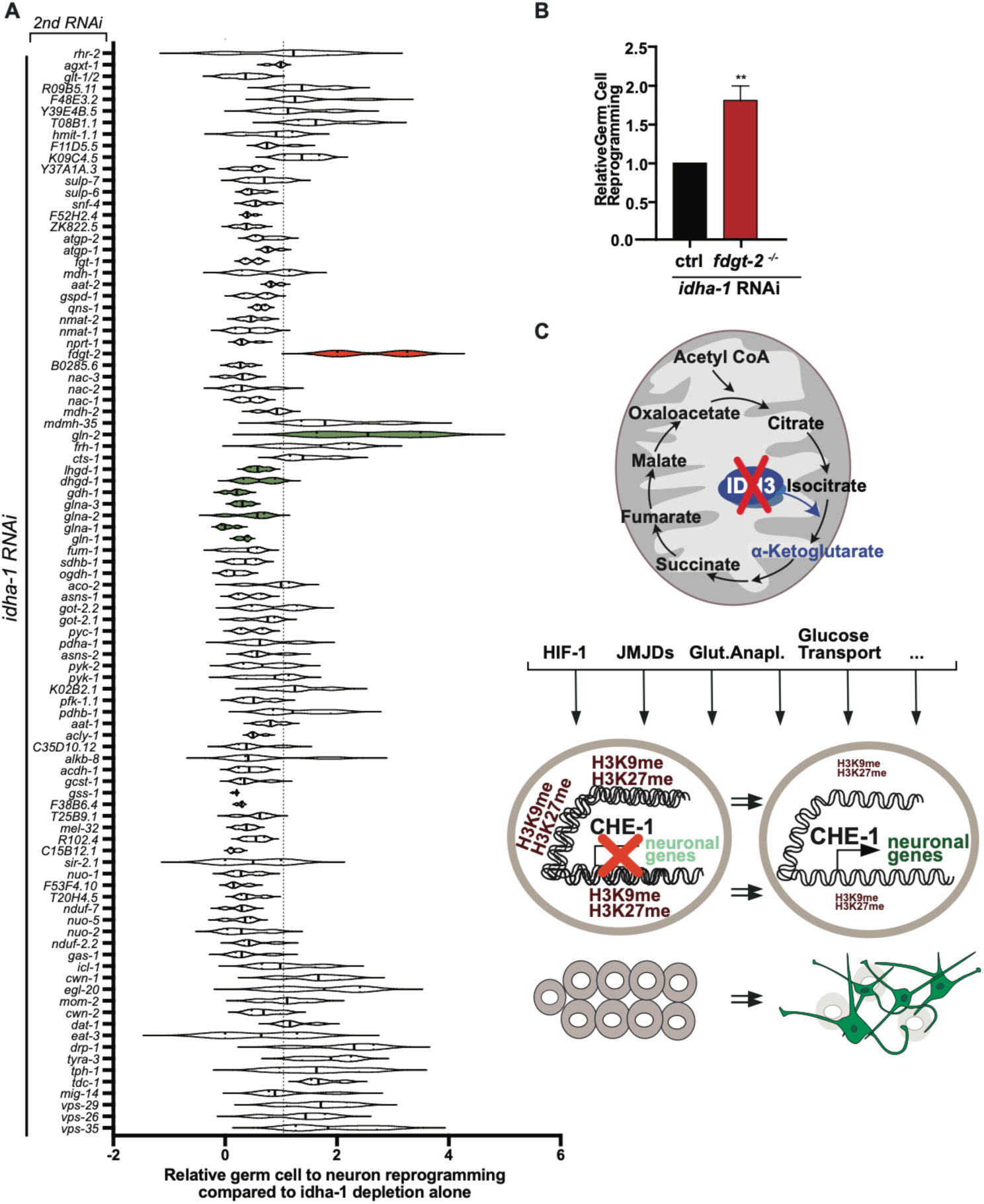
Genetic modifier screening identifies transporter FDGT-2 as a restraint on IDH3 depletion-mediated GeCo. **(A)** Summary of genetic modifiers of IDHA-1 depletion-mediated germ cell reprogramming identified by secondary RNAi screening. Candidate genes were co-depleted with *idha-1*, and CHE-1^OE^- induced GeCo was quantified relative to *idha-1* RNAi alone. Values are normalized to the *idha-1* RNAi condition, indicated by the dashed line at 1; values below or above this reference indicate suppression or enhancement of reprogramming, respectively. The putative monosaccharide transporter *fdgt-2* is highlighted in red, while selected genes involved in glutamine/α-KG metabolism are highlighted in green. The screen identified both suppressors and enhancers of the *idha-1*-dependent phenotype, with depletion of *fdgt-2* producing a prominent enhancement of reprogramming. Data are presented as mean ± SEM. Statistical significance was determined using ordinary one-way ANOVA followed by Dunnett’s multiple-comparisons test, comparing each co-depletion condition with the *idha-1* RNAi control. ***P < 0.001; ns, not significant. **(B)** Genetic validation of FDGT-2 as a negative regulator of IDHA-1 depletion-mediated reprogramming. GeCo following *idha-1* RNAi was quantified in control and *fdgt-2*-/- (CRISPR deletion) animals and normalized to the control response. Loss of *fdgt-2* significantly enhanced reprogramming following IDHA-1 depletion. Data are presented as mean ± SEM. Statistical significance was determined using an unpaired two-tailed Student’s *t*-test. **P < 0.01. **(C)** Integrated model for metabolic regulation of IDH3 depletion-mediated germ cell reprogramming. Loss of mitochondrial IDH3 disrupts the canonical isocitrate-to-α-KG step of the TCA cycle and engages multiple compensatory and signaling responses, including HIF-1 activity, Jumonji-domain histone demethylases, glutamine anaplerosis, and altered monosaccharide transport. HIF-1-dependent signaling, Jumonji activity, and metabolic rerouting toward α-KG are proposed to converge with the observed changes in histone methylation to establish a reprogramming-permissive state. In contrast, FDGT-2-dependent transport provides a compensatory metabolic response that limits the extent of reprogramming. Together, these pathways determine whether differentiated germ cells remain refractory to CHE-1 or acquire competence for neuronal conversion.

This finding indicates that reduced glucose uptake acts in the same direction as elevated HIF-1 and reduced GLN-2, adding a third metabolic input caused by idha-1 depletion that increases germ-cell permissiveness to TF-induced conversion. Together, our results support a model in which perturbation of the TCA cycle at the isocitrate-to-αKG step causes HIF-1 stabilization, JmjC-dependent activities, altered glutamate anaplerosis, and reduced glucose transport, which converge on the germ cell nucleus to diminish the repressive H3K9me3 and H3K27me3 histone modifications, allowing ectopic CHE-1 to activate neuronal gene expression resulting in DR for germ cells into neuron-like cells (Figure 7C).

## DISCUSSION

The most direct interpretation of an IDH3 block causing less αKG, and therefore less activity of αKG-dependent dioxygenases such as JmjC demethylases, resulting in maintained or even more repressive histone methylation is contradicted by our finding. Germline H3K9me3 and H3K27me3 decrease rather than increase, whole-animal steady-state αKG is not measurably reduced by *idha-1* RNAi based on our different metabolomics measurements, and feeding αKG does not suppress conversion. These apparent contradictions are informative rather than merely negative, and they suggest several possible explanations.

We note that GC-MS of whole-worm extracts reports a population average across tissues, and more tissue-resolved information on αKG would provide additional hints towards the causative effects. To test this, we would need to measure the relevant αKG levels for JmjC and PHD enzymes in germ cells by metabolomics on isolated gonads. Our attempt at GC-MS using 100 isolated gonads per sample was unsuccessful, and a direct αKG sensor is currently unavailable. A potential sensor based on the ATP synthase ATP-2 could be an alternative route (Chin, Fu et al. 2014), or collecting more gonads. But establishing these approaches is beyond the scope of the present study. Notably, αKG-dependent dioxygenases are competitively inhibited by succinate and fumarate, so their activity depends on the αKG/succinate and αKG/fumarate ratios rather than on absolute αKG alone (Xiao, Yang et al. 2012, TeSlaa, Chaikovsky et al. 2016). Succinate is modestly reduced at later time points upon *idha-1* RNAi while αKG is unchanged, which would raise the αKG/succinate ratio and thus *increase* dioxygenase activity, consistent with the loss of repressive marks we observe, and with the requirement for JMJD-3.3 and JMJD-4. Yet, the enhancement of GeCo by fumarate and malate feeding does not fit this framework and may instead reflect anaplerotic replenishment of the distal cycle, consistent with the requirement for *ogdh-1*, *sdhb-1,* and *fum-1*. Yet, the requirement for *jmjd-4* is interesting, because the mammalian orthologue JMJD4 is not a histone demethylase. JMJD4 is an αKG-dependent lysyl hydroxylase that hydroxylates the translation termination factor eRF1 at K63 and is required for optimal translational termination (Feng, Yamamoto et al. 2014, Zhuang, Feng et al. 2015). If the *C. elegans* protein retains this activity, JMJD-4 may contribute to GeCo through protein hydroxylation and translational control rather than chromatin.

HIF-1 stabilization by eliminating VHL-1 increases GeCo, while HIF-1 loss decreases it. The fact that several JmjC demethylases are HIF-1 target genes in other systems (Pollard, Loenarz et al. 2008, Wellmann, Bettkober et al. 2008, Krieg, Rankin et al. 2010) provides a plausible feed-forward route by which even a modest change in PHD activity based on αKG levels is translated into a larger change in demethylase dosage. While our study establishes a genetic requirement chain from IDH3 activity to HIF-1, JMJD-3.3/JMJD-4, and permissive chromatin required for GeCo, dissecting the exact metabolic link will require tissue-specific chromatin profiling in germ cells combined with direct measurement of the relevant metabolite pools. These insights are central goals of future work rather than gaps to close in the present study.

Furthermore, we provide evidence that GeCo requires flux through glutamate into αKG upon *idha-1* RNAi, which is supported by metabolomics showing that glutamate is significantly reduced upon *idha-1* RNAi. Glutamine is rather unchanged, indicating increased glutamate consumption rather than a shortage of its precursor, and that the glutamine pool may be buffered by dietary uptake or alternative synthesis routes. Hence, when the canonical route to αKG is blocked at IDH3, cells appear to rely on glutamate anaplerosis as previously shown in other systems (Owen, Kalhan et al. 2002, DeBerardinis, Mancuso et al. 2007, Xiao, Zeng et al. 2016). In particular, the TCA block in cancer cells increases glutamate dehydrogenase (GDH) activity to generate αKG from glutamate, consistent with our finding that GDH-1 is required for GeCo upon TCA block by IDHA-1 depletion (Yang, Ko et al. 2014). Overall, manipulations that increase or decrease the glutamate available for this route in *C. elegans* enhance or suppress DR of germ cells to neurons accordingly.

The targeted RNAi screen for additional metabolic factors identified *fdgt-2*, which encodes a facilitated glucose transporter of the GLUT/SLC2A family (Kim, Underwood et al. 2018), as another strong enhancer. This finding indicated that reduced glucose uptake may limit glycolytic flux and pyruvate supply to the cycle, compounding the IDH3 block. Notably, loss of FDGT-2 at the glucose-uptake step would be expected to increase reliance on glutamate anaplerosis as well (Yang, Ko et al. 2014). While the exact function of FDGT-2 has not yet been studied experimentally, its identification supports the broader point of this study: multiple metabolic perturbations based on impaired αKG production, including HIF-1 stabilization, altered glutamate anaplerosis, and potentially reduced glucose import, shift germ cells towards permissiveness for DR to neuron-like cells.

The list of characterized reprogramming barriers so far is dominated by chromatin factors such as LIN-53/RBBP4-7 (Tursun, Patel et al. 2011), CAF1 (Cheloufi, Elling et al. 2015), PRC2 (Patel, Tursun et al. 2012), FACT (Kolundzic, Ofenbauer et al. 2018) , MRG-1 (Hajduskova, Baytek et al. 2019), and more. Although a metabolic barrier joins this list, epigenetic regulation remains the ultimate mechanism for maintaining germ cell identity through repressive H3K27me3 and H3K9me3 chromatin domains. Our data indicate that this state is metabolically supported and requires IDH3 activity. That conversion proceeds to two neuronal fates (ASE via CHE-1, GABAergic via UNC-30) but not to a muscle fate under the same conditions suggests that removing the metabolic barrier creates sufficient permissiveness only for the neuronal program, potentially because the MyoD homolog HLH-1 alone is not strong enough to induce the muscle fate in the context of reduced metabolic barrier. This result mirrors the fate-selectivity previously observed for chromatin barriers, where FACT depletion permits different conversions in different tissues (Kolundzic, Ofenbauer et al. 2018).

Beyond direct reprogramming, our findings are also related to IDH pathology in humans. Certain IDH1/IDH2 mutations cause excessive production of the oncometabolite 2-hydroxyglutarate, which inhibits αKG-dependent dioxygenases and increases repressive histone methylation, thereby blocking differentiation in glioma and leukemia (Figueroa, Abdel-Wahab et al. 2010, Xu, Yang et al. 2011, Lu, Ward et al. 2012). In the *C. elegans* germline, loss of the NAD-dependent isozyme IDH3 has the opposite consequence since it is required to maintain repressive histone methylation. That two different isocitrate dehydrogenases can push chromatin and cell-fate plasticity in opposite directions underscores how much cell identity is negotiated at this metabolic node of the TCA flux.

## ACKNOWLEDGEMENT

We thank current and former Tursun lab members, including Sergej Herzog, Clara Kraus, Andreas Ofenbauer, Ena Kolundzic, Sefanie Müthel, Marcel Studt, and Taina Averdieck, for research and technical support with *C. elegans* experiments. We thank the CGC, which is funded by the NIH Office of Research Infrastructure Programs (P40 OD010440), for *C. elegans* strains. This work was funded by the Deutsche Forschungsgemeinschaft (DFG Project#: 672623).

## SUPPLEMENTARY FIGURE LEGENDS

**Figure S1.**
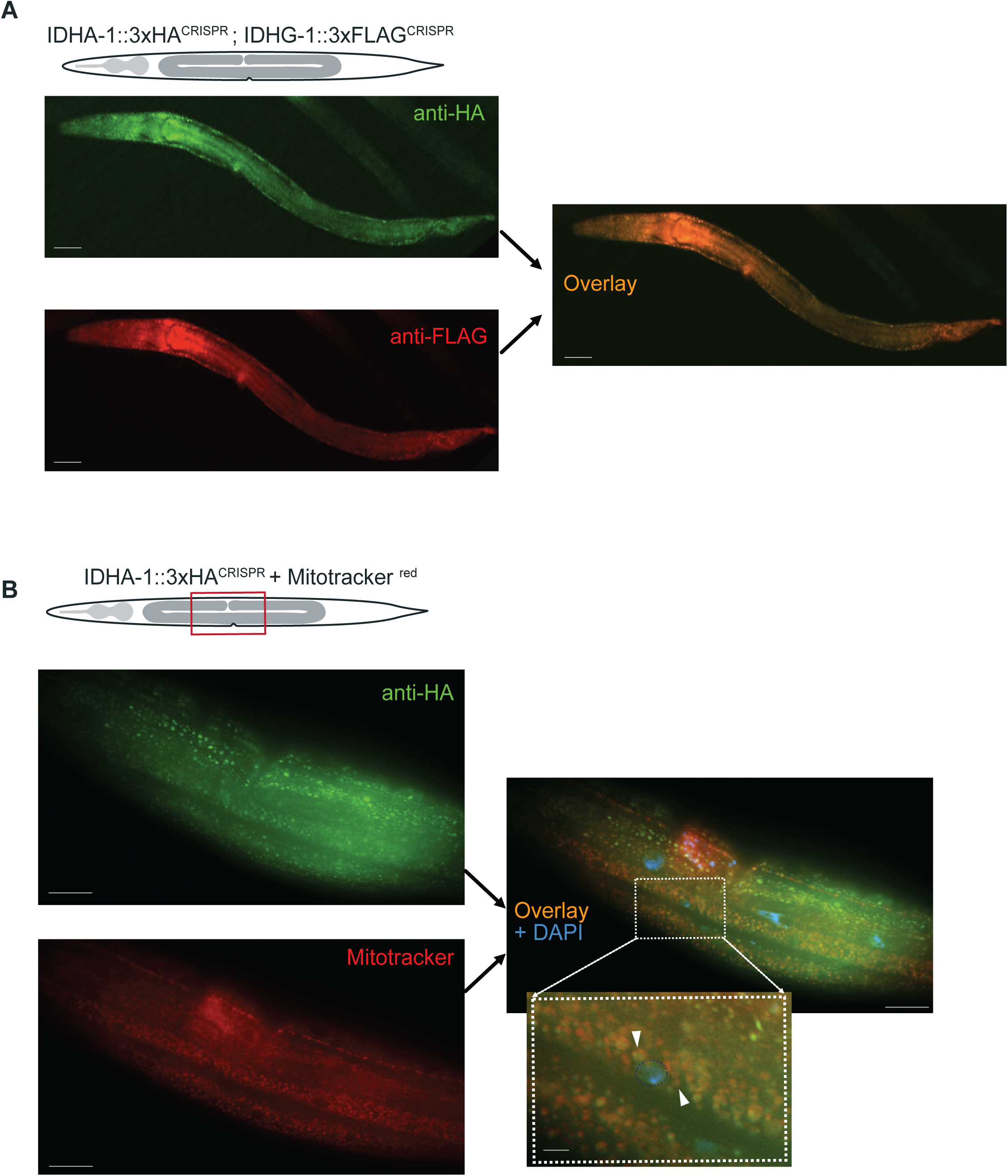
Endogenous IDHA-1 and IDHG-1 show overlapping expression, and IDHA-1 localizes to mitochondria in the *C. elegans* germ line. **(A)** Expression and co-localization of endogenous IDH3 subunits IDHA-1 and IDHG-1. Animals carrying CRISPR-tagged *idha-1::3xHA* and *idhg-1::3xFLAG* alleles were immunostained with anti-HA and anti-FLAG antibodies to visualize the endogenous proteins. IDHA-1::3xHA (green) and IDHG-1::3xFLAG (red) show broad expression throughout the animal and substantial overlap in the merged image, consistent with co-expression of the two IDH3 subunits *in vivo*. Both proteins were tagged at their endogenous loci. **(B)** Mitochondrial localization of endogenous IDHA-1 in the germline region. Animals carrying CRISPR tagged *idha-1::3xHA* were visualized by anti-HA immunofluorescence (green) together with MitoTracker staining (red); nuclei are counterstained with DAPI (blue) in the merged image. The boxed region is shown at higher magnification. IDHA-1 displays a punctate intracellular distribution that overlaps with MitoTracker positive structures, demonstrating mitochondrial localization of IDHA-1 in germ cells. Arrowheads indicate representative regions of overlapping IDHA-1 and mitochondrial signal. Scale bars, 50 μm.

**Figure S2.**
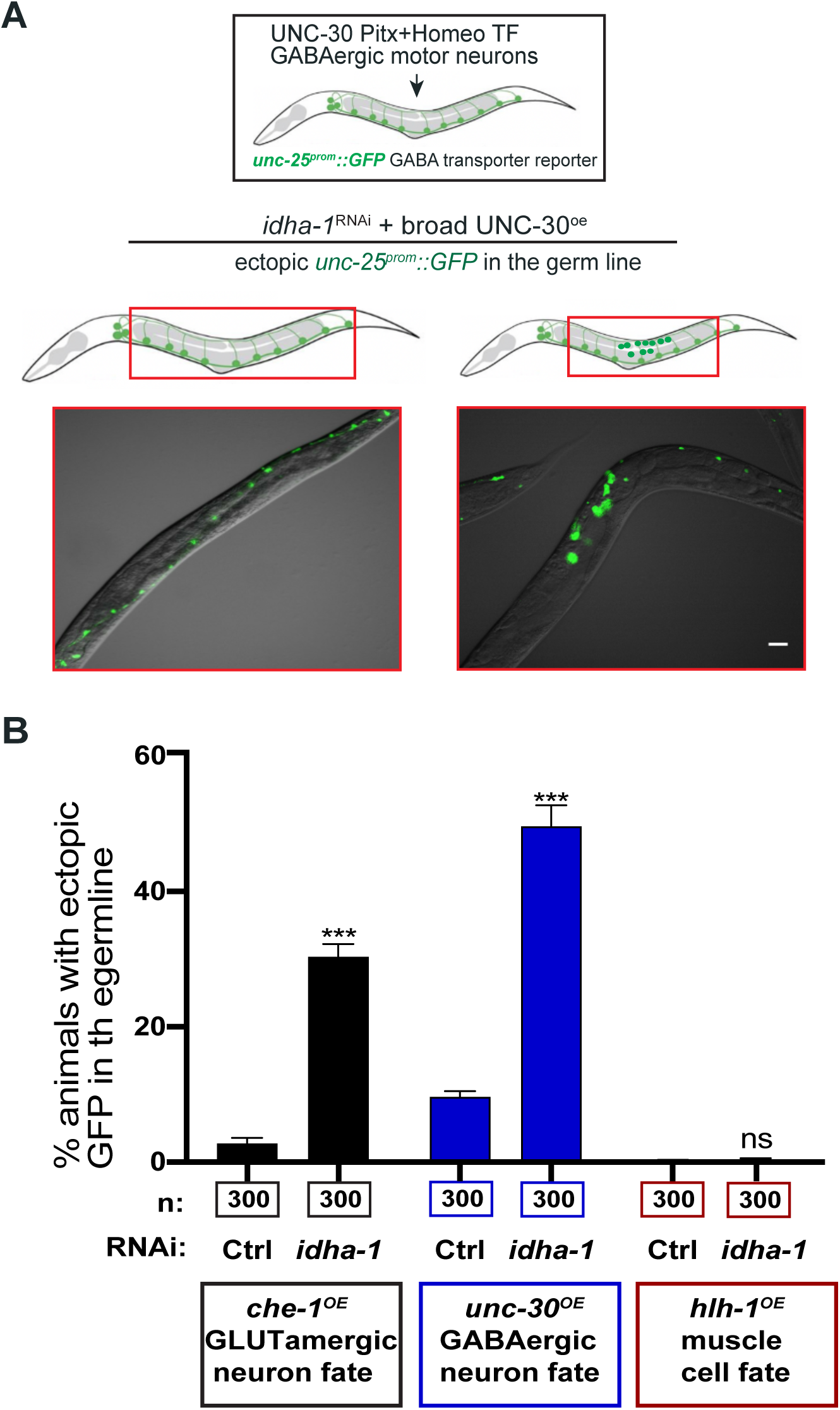
IDHA-1 depletion permits reprogramming toward distinct neuronal fates but not toward muscle fate. **(A)** Assessment of GABAergic neuronal reprogramming following IDHA-1 depletion. UNC-30, a Pitx-type homeodomain transcription factor that specifies GABAergic motor neuron identity, was broadly overexpressed in animals carrying the GABAergic neuronal reporter *unc-25*^prom::^*GFP*. Following *idha-1* RNAi, UNC-30^OE^ induced ectopic *unc-25*^prom^::*GFP* expression in germ cells, whereas reporter expression remained restricted to the endogenous GABAergic neuronal population in control animals. Representative merged DIC/GFP images are shown; boxed regions indicate the germline region. Scale bar, 50 μm. **(B)** Comparison of germ cell reprogramming toward glutamatergic neuronal, GABAergic neuronal, and muscle fates following *idha-1* depletion. Broad overexpression of CHE-1 or UNC-30 was used to induce glutamatergic ASE neuronal or GABAergic neuronal identity, respectively, whereas HLH-1, the *C. elegans* MyoD homolog, was used to induce muscle fate. Following *idha-1* RNAi, CHE-1^OE^ and UNC-30^OE^ produced significant ectopic activation of the corresponding neuronal fate reporters in the germline. In contrast, HLH-1^OE^ failed to induce ectopic muscle reporter expression after idha-1 depletion. These data suggest that loss of IDHA-1 creates a permissive state for neuronal reprogramming rather than a general loss of germ cell fate restriction. Numbers below the bars indicate the total number of animals scored. Data are presented as mean ± SEM. Statistical significance was determined by ordinary one-way ANOVA followed by Sidak’s multiple-comparisons test. ***P < 0.001; ns, not significant.

**Figure S3.**
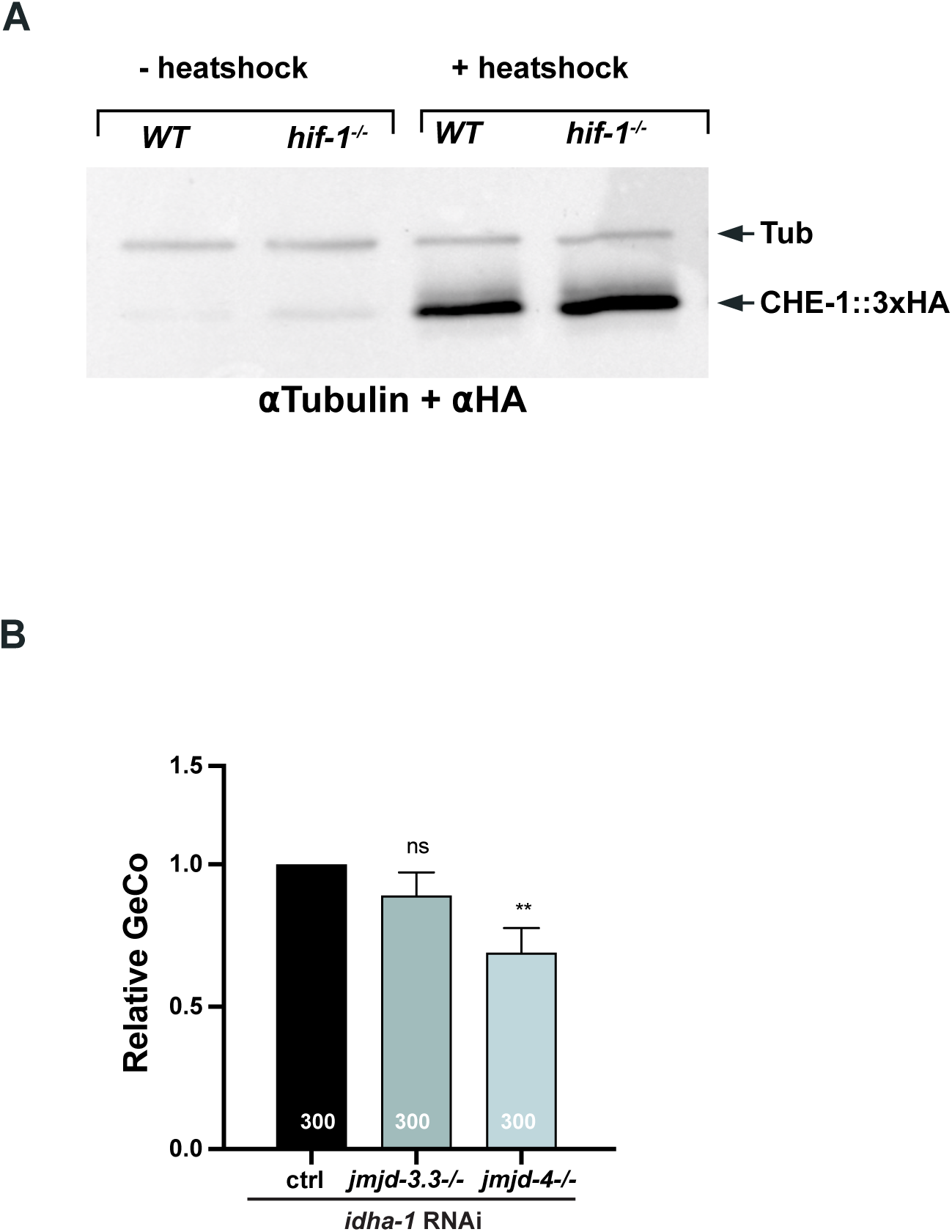
HIF-1 loss does not impair CHE-1 induction, and genetic disruption of *jmjd-4* suppresses IDHA-1 depletion-mediated reprogramming. **(A)** HIF-1 loss does not alter heat-shock-induced CHE-1 expression. Whole animal lysates from wild-type and *hif-1*-/- animals carrying the heat-shock inducible CHE-1::3xHA transgene were analysed before and after heat shock by western blotting using anti HA antibodies. CHE-1::3xHA was undetectable or present only at background levels in the absence of heat shock and was robustly induced following heat shock in both wild-type and *hif-1*-/- animals. Comparable CHE-1::3xHA levels following induction indicate that suppression of IDHA-1 depletion-mediated reprogramming in the *hif-1*mutant background is not attributable to reduced CHE-1 expression. α-tubulin was used as a loading control. **(B)** Genetic validation of candidate Jumonji factors identified in the RNAi suppressor screen. CHE-1^OE^ induced GeCo following *idha-1* RNAi was quantified in control, *jmjd-3.3*-/-, and *jmjd-4*-/- animals and normalized to the control response. Loss of *jmjd-4* significantly suppressed IDHA-1 depletion mediated reprogramming, whereas loss of *jmjd-3.3* did not significantly alter GeCo. These results independently support a requirement for JMJD-4 in the reprogramming response and are consistent with the suppression observed following *jmjd-4* RNAi. Numbers within bars indicate the total number of animals scored. Data are presented as mean ± SEM. Statistical significance was determined by one-way ANOVA followed by Dunnett’s multiple-comparisons test, comparing each mutant with the control. **P < 0.01; ns, not significant.

**Figure S4.**
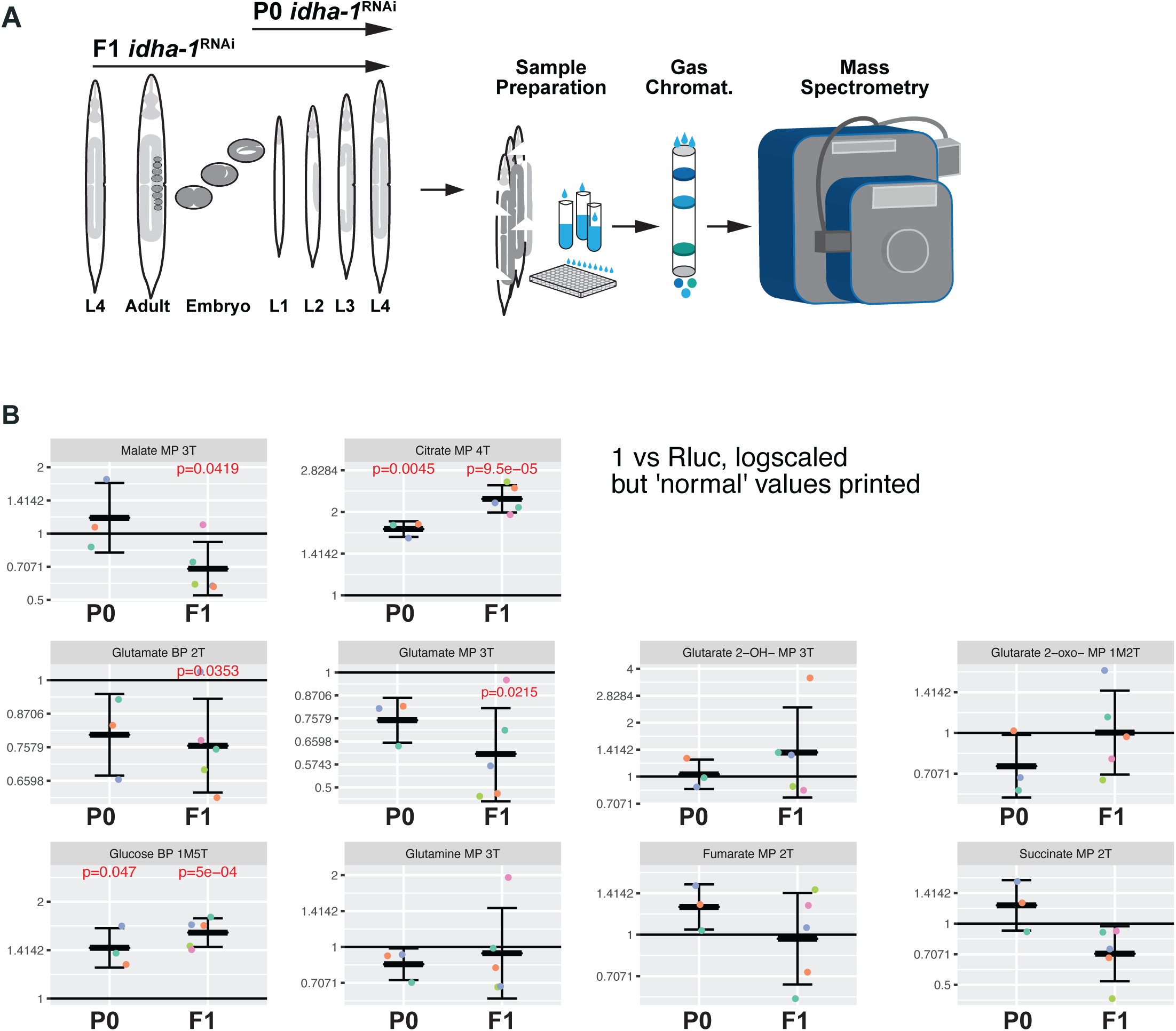
IDH3 depletion alters metabolite abundance while maintaining downstream TCA cycle intermediates. **(A)** Schematic of the GC-MS-based metabolomics workflow. Animals were subjected to control or *idha-1* RNAi and collected at the L4 stage following P0 or F1 RNAi treatment. Metabolites were extracted from whole animals, derivatized, and analyzed by gas chromatography-mass spectrometry (GC-MS). Metabolite abundance was quantified from corrected chromatographic peak areas and expressed relative to the corresponding control condition. **(B)** Relative abundance of selected TCA cycle and associated metabolites following *idha-1* depletion. Values represent fold change in *idha-1* RNAi-treated animals relative to the corresponding *Rluc* RNAi control, with a value of 1 indicating no change. Individual coloured points represent independent biological replicates. Citrate was markedly increased following *idha-1* depletion in both P0 and F1 animals, consistent with accumulation of metabolites upstream of the disrupted IDH3 node. In contrast, downstream TCA cycle intermediates, including fumarate and succinate, showed no consistent reduction, while malate was altered. α-KG was also not significantly reduced in the original metabolomic analysis, supporting maintenance of the downstream TCA cycle metabolite pool despite loss of IDH3. Glutamate and related metabolites were additionally examined because glutamate can provide an alternative route to α-KG; reduced glutamate abundance following *idha-1* depletion is consistent with altered glutamate metabolism in this context.

**Figure S5.**
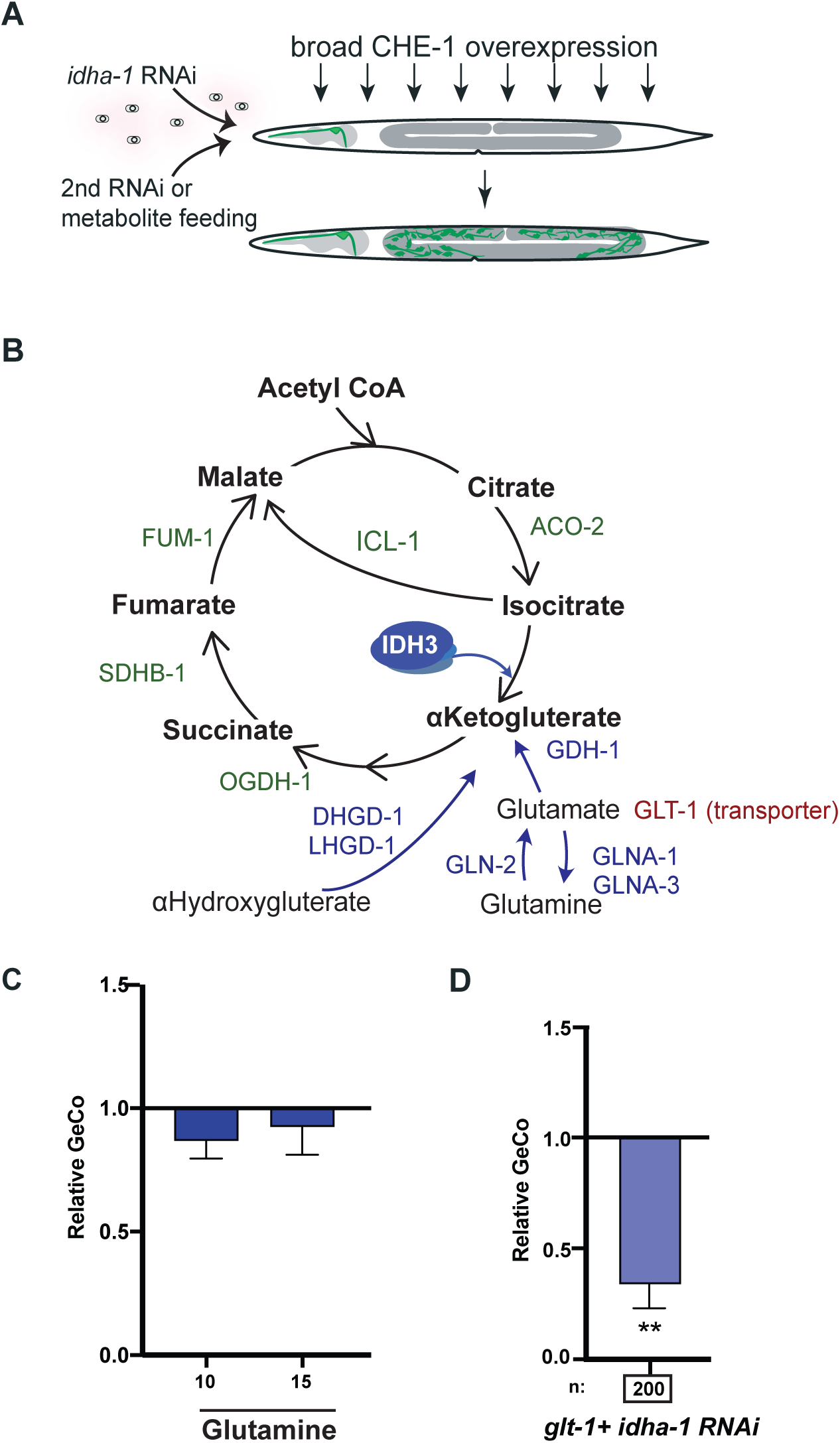
Glutamine supplementation does not further enhance reprogramming, whereas glutamate transport contributes to the IDH3 depletion mediated phenotype. **(A)** Schematic of the experimental strategy used to test metabolic modifiers of IDHA-1 depletion-mediated germ cell reprogramming. Animals were treated with *idha-1* RNAi together with either a second RNAi perturbation or exogenous metabolite supplementation, followed by broad CHE-1 overexpression. GeCo was assessed by ectopic activation of the neuronal fate reporter in the germline. **(B)** Schematic of α-KG generation and downstream TCA cycle pathways examined genetically or by metabolite supplementation. **(C)** Effect of exogenous glutamine supplementation on IDHA-1 depletion-mediated reprogramming. Animals treated with *idha-1* RNAi were supplemented with glutamine at the indicated concentrations, and GeCo penetrance following CHE-1^OE^ was quantified relative to the untreated *idha-1* RNAi condition. Glutamine supplementation did not significantly alter reprogramming, indicating that increasing extracellular glutamine availability is not sufficient to further enhance the IDH3 depletion-mediated phenotype. Data are presented as mean ± SEM from three independent biological experiments. Statistical significance was determined by one-way ANOVA followed by Dunnett’s multiple comparisons test, comparing each glutamine concentration with the *idha-1* RNAi control; ns, not significant. **(D)** Requirement for the glutamate transporter GLT-1 during IDHA-1 depletion-mediated reprogramming. Co-depletion of *glt-1* with *idha-1* markedly reduced GeCo relative to *idha-1* RNAi alone, supporting a functional contribution of glutamate transport to the metabolic state required for reprogramming. The result is consistent with glutamate availability or transport being important for the glutamine/glutamate to α-KG pathway. Numbers below the bars indicate the total number of animals scored. **Statistical significance was determined using a two-tailed one-sample Student’s *t*-test. P < 0.01.

**Figure S6.**
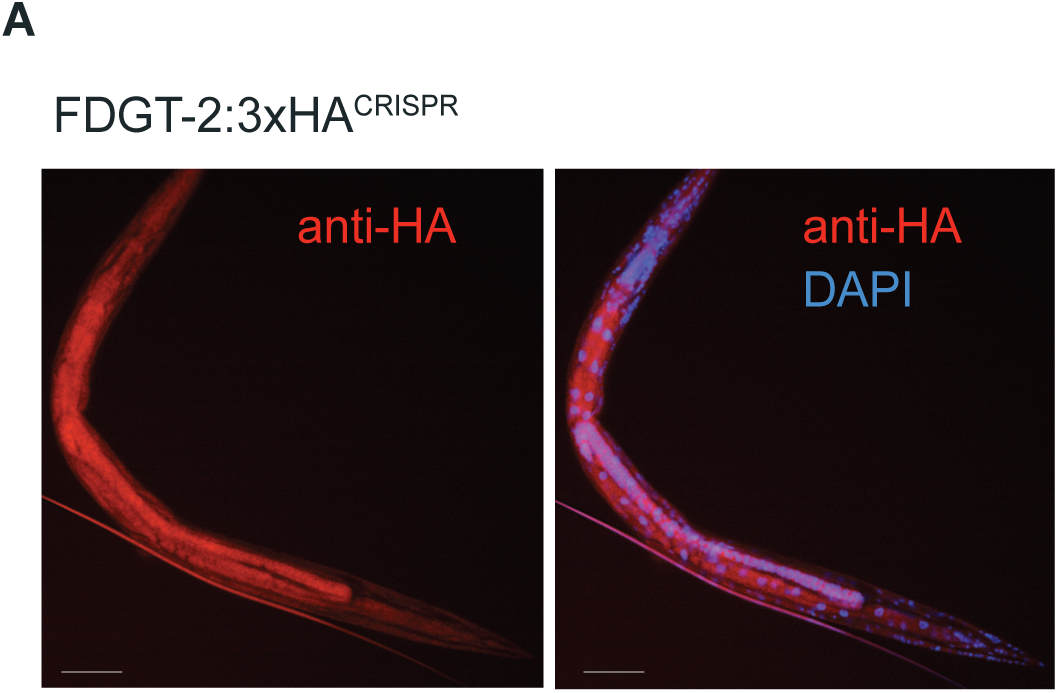
FDGT-2 is broadly expressed in *C. elegans*. **(A)** Representative immunofluorescence images of animals carrying a CRISPR-engineered endogenous *fdgt-2::3xHA* allele. FDGT-2::3xHA was detected by anti-HA immunostaining (red), and nuclei were counterstained with DAPI (blue). FDGT-2 shows broad expression throughout the animal. The merged image shows FDGT-2::3xHA together with nuclear

## MATERIAL AND METHODS

### *C. elegans* Strains Used in the Study

Information on strains, including genotypes, is provided in Table S1. Hermaphrodites were used for phenotype scoring including reprogramming assays, reporter analysis, smFISH, immunohistochemistry and biochemical experiments. Endogenously tagged and mutant strains generated in this study are described below.

### Nematode Culture

*C. elegans* strains were maintained using standard conditions on OP50 bacteria (Brenner, 1974). All heat-shock and temperature-sensitive strains were maintained at 15°C. Animals were synchronized by sodium hypochlorite treatment of gravid adults and embryos were washed with M9 buffer before transfer to NGM plates.

### RNAi in C. elegans

For RNAi, worms were grown on NGM plates containing 1 mM IPTG and 50 μg/ml carbenicillin and seeded with RNAi bacteria from the Ahringer library (Source BioScience). RNAi against *Renilla luciferase* (Rluc) was used as control. Reprogramming experiments were carried out using a standard RNAi feeding protocol (Kamath et al., 2003). For P0 experiments, worms were synchronized by bleaching and embryos were placed directly on RNAi plates. For F1 RNAi, synchronized animals were grown at 15°C on normal food until the L4 stage and transferred to RNAi plates. Worms were maintained at 15°C until most P0 or F1 progeny reached the L4 stage. Unless stated otherwise, reprogramming experiments were performed using F1 RNAi.

To induce broad CHE-1 expression, plates were heat-shocked at 37°C for 30 min followed by overnight incubation at 25°C as described previously (Kolundzic et al., 2018; Tursun et al., 2011). Animals were screened for ectopic *gcy-5prom::GFP* expression in the germline the following day using a fluorescence dissecting microscope. Germ cell conversion (GeCo) was scored as the percentage of animals displaying ectopic reporter expression in the germline. This assay was used to assess IDH3 subunits, related isocitrate dehydrogenases and genetic modifiers of IDHA-1 depletion-mediated reprogramming.

For double RNAi experiments, saturated bacterial cultures were mixed in a 1:1 ratio based on OD600 and subsequently seeded on RNAi plates. For co-depletion experiments with *idha-1*, bacteria expressing *idha-1* dsRNA mixed with Rluc RNAi bacteria were used as the corresponding control. Candidate genes involved in chromatin regulation, TCA-cycle metabolism, glutamine metabolism and other metabolic pathways were tested for suppression or enhancement of the *idha-1* depletion-mediated GeCo phenotype.

For selected genetic interaction experiments, stitched RNAi constructs were generated to simultaneously target *idha-1*together with *jmjd-3.3* or *jmjd-4*. Target sequences were amplified from existing RNAi plasmids and inserted into an RNAi vector containing the second target sequence by Gibson Assembly. Constructs were confirmed by Sanger sequencing and transformed into HT115(DE3) bacteria. An *idha-1::Rluc* construct was used as control.

### Transcription Factor-Induced Reprogramming

CHE-1-mediated conversion of germ cells toward ASE neuronal identity was assessed using animals carrying heat-shock-inducible *che-1* together with the ASE fate reporter *gcy-5prom::GFP*. To determine whether IDHA-1 depletion also permits conversion toward an alternative neuronal identity, UNC-30 was broadly induced in animals carrying the GABAergic neuronal reporter *unc-25prom::GFP*. Reprogramming toward muscle fate was tested by broad induction of the myogenic transcription factor HLH-1 in animals carrying the muscle reporter *unc-97::GFP*. Reporter induction in germ cells was assessed following control or *idha-1*RNAi.

### Generation of CRISPR Alleles

CRISPR engineering was performed by microinjection using sgRNAs and recombinant Cas9 protein as described previously (Dokshin et al., 2018). For knock-in alleles, repair templates carrying homology arms were also included. A plasmid expressing *myo-2::RFP* was included as a co-injection marker. F1 progeny displaying pharyngeal RFP were screened by PCR for the desired modification. Positive animals were homozygosed by singling and the edited locus was confirmed by Sanger sequencing.

This approach was used to generate endogenously tagged *idha-1::3xHA* and *idhg-1::3xFLAG* alleles and mutant alleles of candidate genes identified in genetic screens. sgRNA, repair templates and genotyping primer sequences are provided in Table S2A-S2C.

### Antibody Staining

For detection of FLAG and UNC-10/RIM in whole worms, animals were fixed and permeabilized as described previously (Bettinger et al., 1996). In brief, animals were washed with M9 and resuspended in RFB (160 mM KCl, 40 mM NaCl, 20 mM EGTA, 10 mM spermidine) supplemented with 2% formaldehyde followed by three freeze-thaw cycles. After incubation for 30 min at 25°C, samples were washed with TTE (100 mM Tris pH 7.4, 1% Triton X-100, 1 mM EDTA) and incubated for 4 h at 37°C in TTE containing 1% β-mercaptoethanol. Samples were washed with BO3 buffer (10 mM H3BO3, 10 mM NaOH, 2% Triton X-100), incubated for 15 min at 37°C in BO3 containing 10 mM DTT, followed by incubation for 15 min at 25°C in BO3 containing 0.3% H2O2. Samples were blocked in PBS containing 0.2% gelatin and 0.25% Triton X-100 for 1 h.

Primary antibodies were diluted in PBS containing 0.25% Triton X-100 and 0.2% gelatin and incubated with samples overnight at 4°C. After washing, Alexa Fluor-conjugated secondary antibodies were applied at 1:1500 and incubated overnight at 4°C. Samples were washed and mounted on glass slides using DAPI-containing mounting medium (Dianova).

For anti-HA, anti-H3K4me3, anti-H3K9me3 and anti-H3K27me3 staining, worms were processed using the slide-crack method as described previously (Jones et al., 1996). Briefly, animals suspended in M9 were placed between two glass slides on dry ice and the slides were separated after freezing. Samples were fixed for 10 min in 4% paraformaldehyde for histone modification staining or methanol for HA staining, washed three times in PBST (0.1% Tween-20 in PBS), blocked for 1 h in 0.2% gelatin in PBST and stained with primary and secondary antibodies as described above. Antibodies used in this study are listed in the Key Resources Table.

### MitoTracker Staining

Mitochondria were labelled using MitoTracker Red CMXRos as described previously (Sarasija and Norman, 2015, 2018). Animals were washed from OP50-seeded plates and incubated in M9 containing 1 μg/ml MitoTracker Red CMXRos for 6 h at 20°C in the dark. Animals were subsequently washed four times with M9, transferred to OP50-seeded NGM plates and allowed to destain overnight at 20°C in the dark. MitoTracker-labelled animals carrying the endogenous *idha-1::3xHA* allele were subsequently fixed and processed for anti-HA immunofluorescence as described above.

### Single Molecule Fluorescent *In Situ* Hybridization (smFISH)

smFISH was performed using Custom Stellaris FISH probes purchased from Biosearch Technologies according to the manufacturer’s protocol. Briefly, animals treated with control or *idha-1* RNAi were washed from plates with M9 followed by five washes with nuclease-free water and fixed in 4% paraformaldehyde for 45 min. Samples were washed twice with DEPC-treated PBS and permeabilized in 70% ethanol overnight at 4°C.

Animals were washed with buffer containing formamide, SSC and Triton X-100 and incubated with the respective probe set in hybridization buffer for 16 h at 37°C in the dark. Samples were washed at 37°C, counterstained with DAPI and mounted using Vectashield mounting medium. Probe sets against endogenous neuronal genes including *gcy-5, rab-3, unc-119* and *unc-10* were used to assess acquisition of neuronal gene expression following IDHA-1 depletion and CHE-1 induction. Sequences of all smFISH probes used are provided in Table S3.

### Fluorescence Microscopy

Animals were imaged using a Zeiss Axio Imager 2 fluorescence microscope equipped with a Sensicam digital camera (PCO) or a Leica DM6B fluorescence microscope. Live animals were mounted on 2% agarose pads in M9 containing 20 mM tetramisole for immobilization.

For quantification of histone modifications, fluorescence intensities within the germline were measured from animals stained for H3K4me3, H3K9me3 or H3K27me3 and compared between control and *idha-1* RNAi conditions. HTZ-1/H2A.Z staining was used as a chromatin reference.

### Western Blotting

For preparation of whole-worm protein lysates, animals were collected at the L4 stage following RNAi treatment and washed three times with M9 to remove bacteria. Worm pellets were resuspended in 5× SDS sample buffer containing 10% SDS, 10 mM DTT, 20% glycerol, 0.2 M Tris-HCl pH 6.8 and 0.05% bromophenol blue and incubated at 96°C for 10 min. Protein samples were separated on 4–20% Mini-PROTEAN precast gels (Bio-Rad) and transferred to nitrocellulose membranes by wet transfer at 100 V for 60 min. Membranes were blocked in 3% BSA in TBST for 1 h at room temperature and incubated with primary antibodies for 4 h at room temperature or overnight at 4°C. After washing, HRP-conjugated secondary antibodies were applied for 1 h at room temperature. Signals were detected using Lumi-Light Western Blotting substrate and an ImageQuant LAS4000 system.

Western blotting was used to analyze IDHA-1 depletion, histone modifications H3K4me3, H3K9me3, and H3K27me3, and heat-shock-induced CHE-1::3xHA expression. α-tubulin was used as loading control.

### Metabolite Feeding

For metabolite supplementation experiments, 400 mM stock solutions were prepared and diluted into NGM agar. The pH of the medium was adjusted to 6 as described previously (Chin et al., 2014). Plates were seeded with RNAi bacteria and animals were subjected to F1 RNAi as described above.

Animals treated with *idha-1* RNAi were supplemented with α-ketoglutarate, succinate, fumarate, malate or citrate at the indicated concentrations and GeCo penetrance was assessed following CHE-1 induction. Glutamine supplementation was performed similarly using the concentrations indicated in the corresponding figure.

### GC-MS Metabolomic Analysis

Metabolomic profiling was performed by gas chromatography coupled to mass spectrometry (GC-MS) in collaboration with the Kempa laboratory at the Berlin Institute for Medical Systems Biology. Animals carrying endogenously tagged *idha-1::3xHA* were subjected to P0 or F1 RNAi. Animals were collected at the L4 stage, washed three times with M9 to remove residual bacteria, and the worm pellets were weighed and immediately flash frozen in liquid nitrogen. Up to 50 mg of worm material was used for metabolite extraction.

Metabolites were extracted by addition of methanol:chloroform:water (5:2:1; MCW) at 1 mL per 50 mg of worm pellet. Samples were transferred to tubes containing silica beads and mechanically disrupted using a tissue lyser at 6.5 m/s with two 20 s pulses separated by 5 s, repeated three times, followed by sonication for 10 min in an ultrasound bath. The supernatant was collected and the remaining extraction solvent was added, followed by vortexing and brief incubation on dry ice. Samples were shaken for 15 min and centrifuged at 1,400 rpm at 4°C. Water corresponding to 0.5 volumes of the extraction mixture was then added for phase separation. Samples were vortexed and shaken for a further 15 min at 1,400 rpm at 4°C, followed by centrifugation at 20,000 × *g* for 10 min at 4°C. The upper polar phase was collected and dried overnight in a vacuum concentrator.

Dried metabolites were derivatized using a modified protocol based on Roessner-Tunali et al. (2003), as described in Kempa et al. (2009). Briefly, 10 μL of 40 mg/mL methoxyamine hydrochloride in pyridine was added, and samples were incubated for 90 min at 30°C with shaking. Subsequently, 30 μL *N*-methyl-*N*-(trimethylsilyl) trifluoroacetamide (MSTFA), supplemented with alkanes as retention-index standards, was added and samples were incubated for 60 min at 37°C. Samples were centrifuged at 20,000 × *g* for 10 min at room temperature and transferred to glass vials for GC-MS analysis.

Identification mixes were included during sample processing and quantification mixes were used at serial dilutions ranging from 1:1 to 1:200 as described previously (Pietzke et al., 2014). GC-MS measurements were performed by Jenny Grobe and Tobias Opialla using procedures established in the Kempa laboratory (Pietzke et al., 2014). Data analysis was performed by Tobias Opialla using Maui-SILVIA (Kuich et al., 2014).

### Stable-Isotope Tracing with ^13^C-Glucose

Stable-isotope tracing was performed separately from the metabolite-abundance analysis to assess incorporation of glucose-derived carbon into downstream metabolic intermediates. Synchronized N2 animals were subjected to F1 control (Rluc) or *idha-1* RNAi as described above and grown to the L3/L4 stage. Animals were then collected in M9 and washed to remove residual bacteria.

Animals were starved in M9 for 1 h before isotope feeding. In parallel, OP50 bacteria were UV-inactivated to prevent bacterial metabolism of the labeled substrate. UV-killed bacteria were collected in ddH₂O and supplemented with either 25 mM ^13^C glucose or 25 mM unlabelled ^12^C-glucose as the corresponding control. Following starvation, animals were resuspended in the respective bacterial/glucose suspensions and collected after 60, 180, 360, and 840 min of isotope labeling. For sample collection, animals were transferred into pre-weighed 1.5 mL tubes and washed with ice-cold M9 until residual bacteria were removed. Worm pellets were weighed, immediately flash-frozen in liquid nitrogen, and stored at -80°C until metabolite extraction. Approximately 30-45 mg of worm material was collected per sample.

Frozen samples were subsequently subjected to GC-MS-based metabolite extraction and analysis. The ^13^C-labelled fraction of individual metabolites was determined by GC-MS and compared between control and *idha-1* RNAi conditions across the labelling time course. Stable isotope labelling was used to trace carbon utilization through metabolic pathways as described previously (Pietzke et al., 2014; Antoniewicz, 2018).

### Quantification and Statistical Analysis

Statistical analyses were performed using GraphPad Prism. Applied statistical tests are indicated in the respective figure legends. Data are presented as mean ± SEM unless otherwise stated. Ordinary one-way ANOVA followed by Dunnett’s, Tukey’s or Sidak’s multiple-comparisons tests was used as indicated. Student’s *t*-tests were used for two-group or normalized single reference comparisons as specified in the corresponding figure legends. *P* < 0.05 was considered statistically significant.

---

**Table S1.**
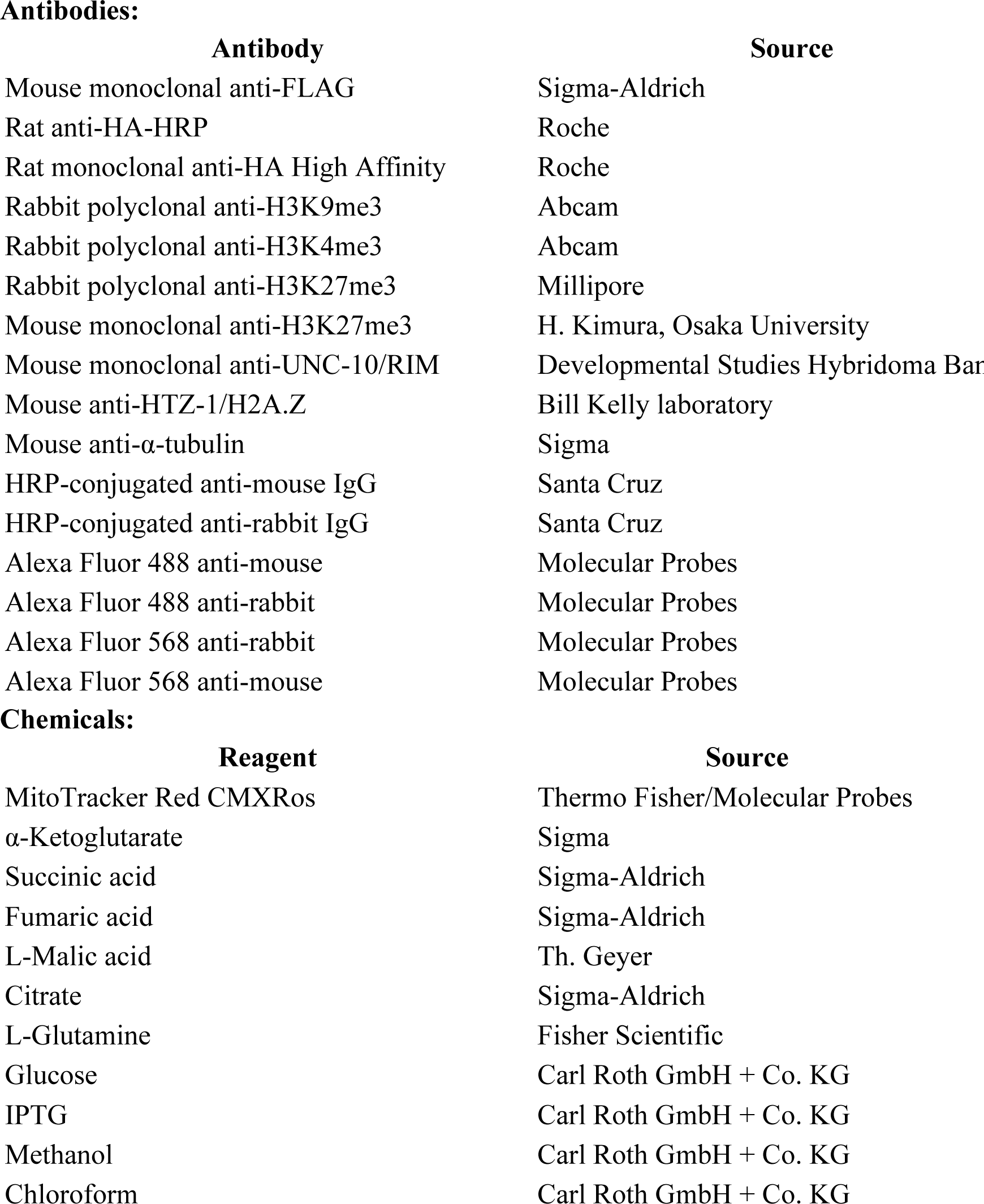
*C. elegans* strains used in this study:

**Antibodies:**
| **Antibody** | **Source** |
| --- | --- |
| Mouse monoclonal anti-FLAG | Sigma-Aldrich |
| Rat anti-HA-HRP | Roche |
| Rat monoclonal anti-HA High Affinity | Roche |
| Rabbit polyclonal anti-H3K9me3 | Abcam |
| Rabbit polyclonal anti-H3K4me3 | Abcam |
| Rabbit polyclonal anti-H3K27me3 | Millipore |
| Mouse monoclonal anti-H3K27me3 | H. Kimura, Osaka University |
| Mouse monoclonal anti-UNC-10/RIM | Developmental Studies Hybridoma Bank |
| Mouse anti-HTZ-1/H2A.Z | Bill Kelly laboratory |
| Mouse anti-α-tubulin | Sigma |
| HRP-conjugated anti-mouse IgG | Santa Cruz |
| HRP-conjugated anti-rabbit IgG | Santa Cruz |
| Alexa Fluor 488 anti-mouse | Molecular Probes |
| Alexa Fluor 488 anti-rabbit | Molecular Probes |
| Alexa Fluor 568 anti-rabbit | Molecular Probes |
| Alexa Fluor 568 anti-mouse | Molecular Probes |

**Chemicals:**
| **Reagent** | **Source** |
| --- | --- |
| MitoTracker Red CMXRos | Thermo Fisher/Molecular Probes |
| α-Ketoglutarate | Sigma |
| Succinic acid | Sigma-Aldrich |
| Fumaric acid | Sigma-Aldrich |
| L-Malic acid | Th. Geyer |
| Citrate | Sigma-Aldrich |
| L-Glutamine | Fisher Scientific |
| Glucose | Carl Roth GmbH + Co. KG |
| IPTG | Carl Roth GmbH + Co. KG |
| Methanol | Carl Roth GmbH + Co. KG |
| Chloroform | Carl Roth GmbH + Co. KG |

**Table S2.**
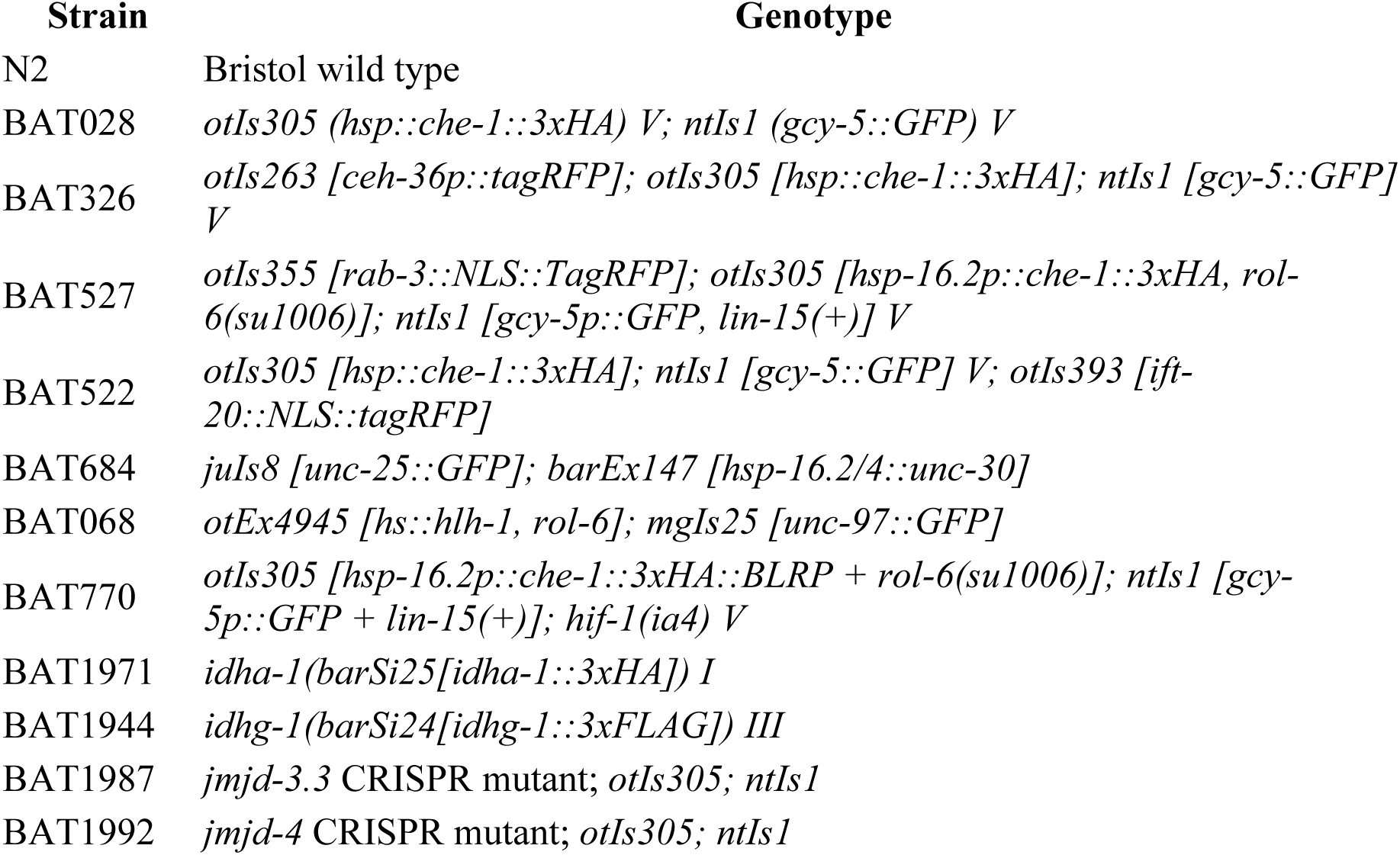
CRISPR reagents and genotyping primers.

**Table S2A.**
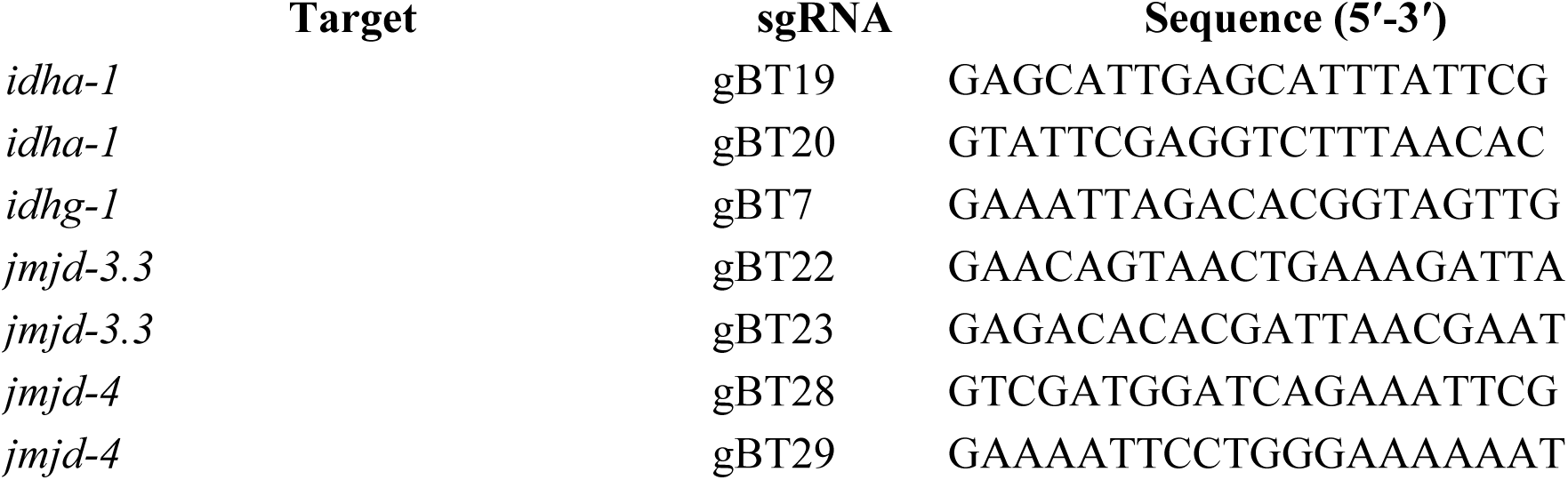
sgRNAs.

**Table S2B.**
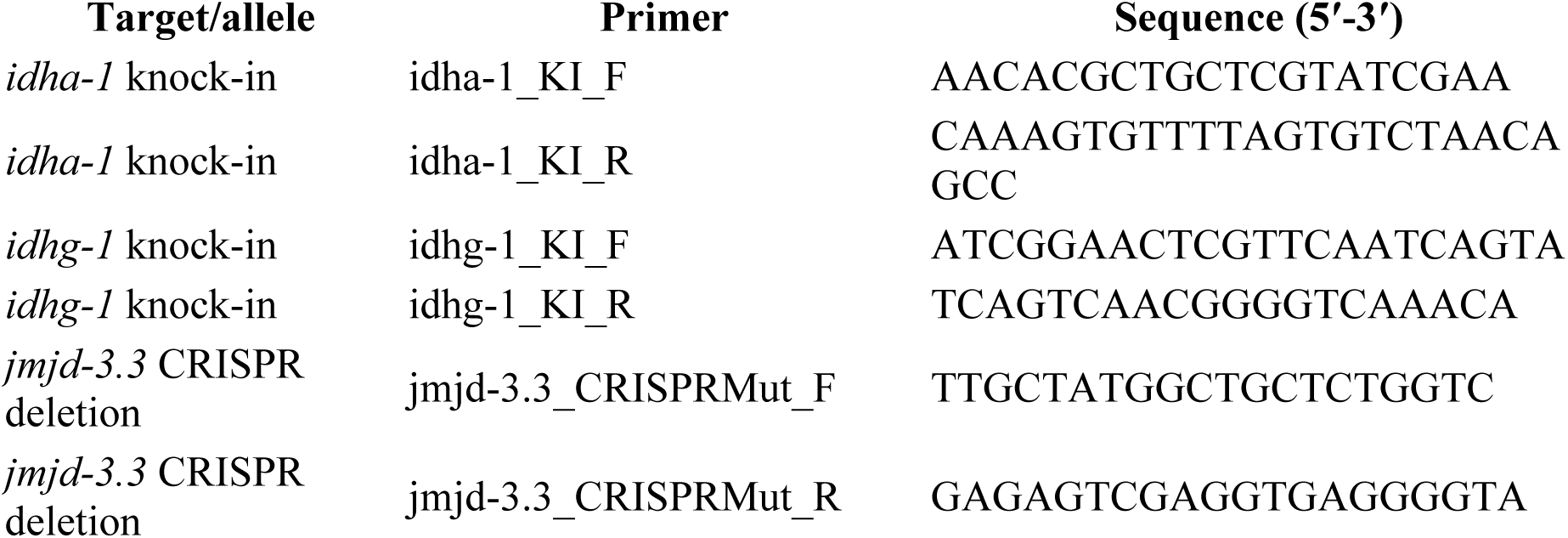
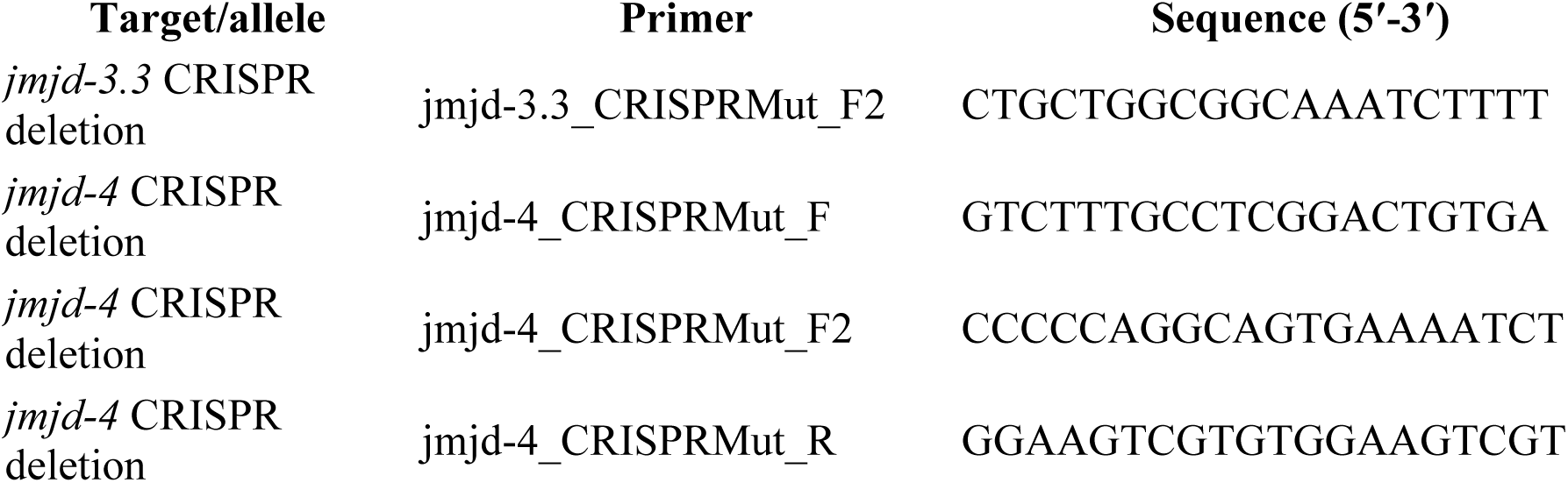
Genotyping primers.

**Table S2C.**
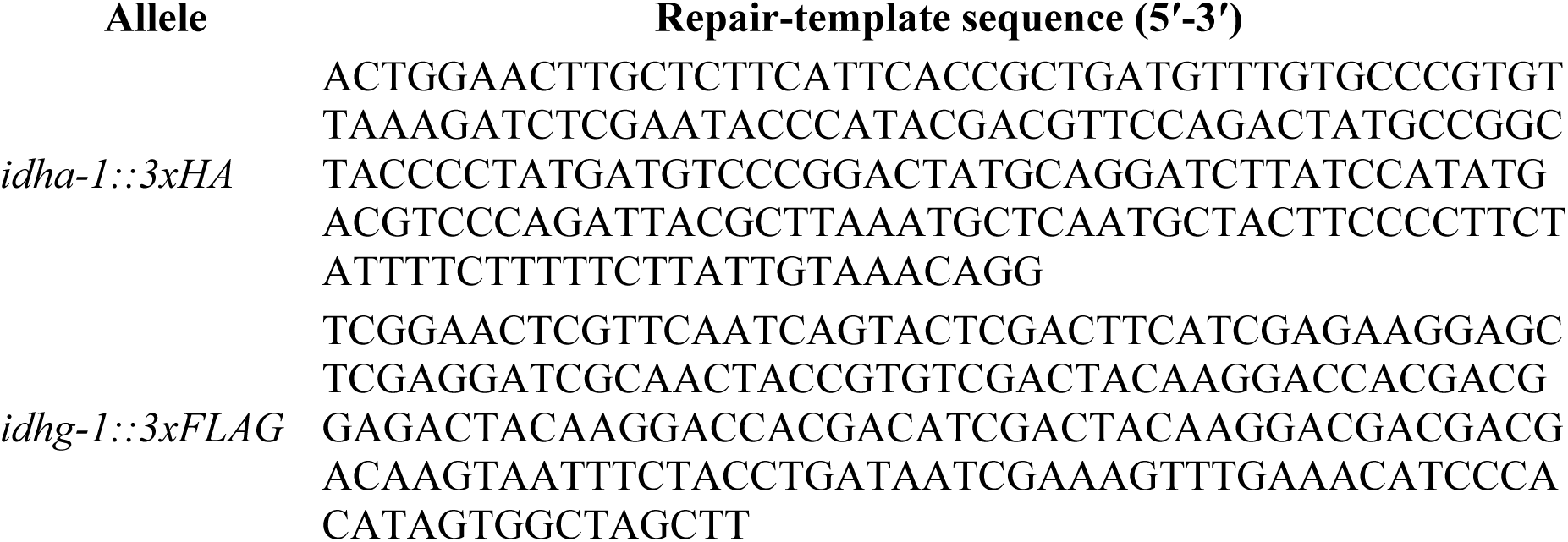
Repair templates.

**Table S3.**
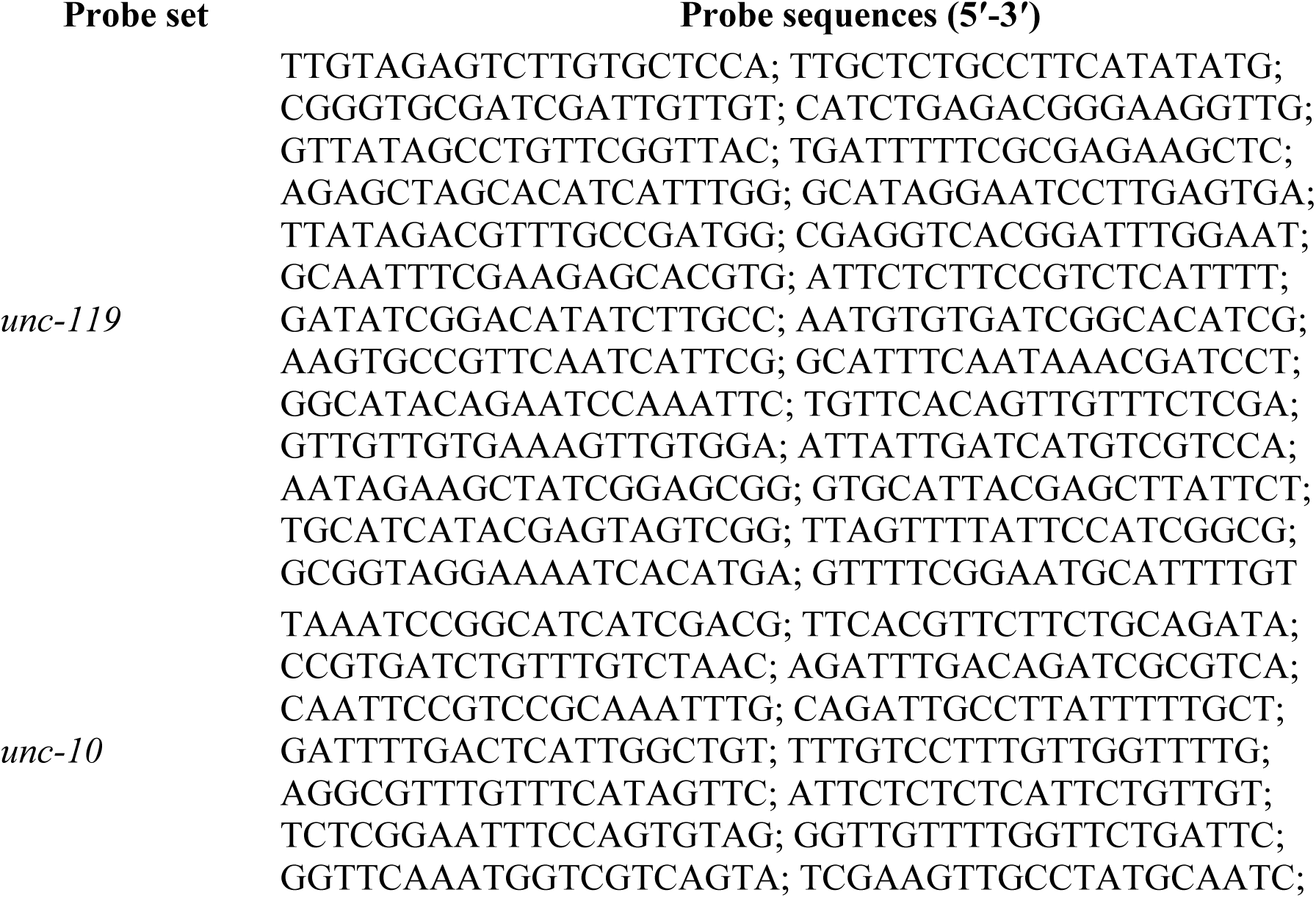
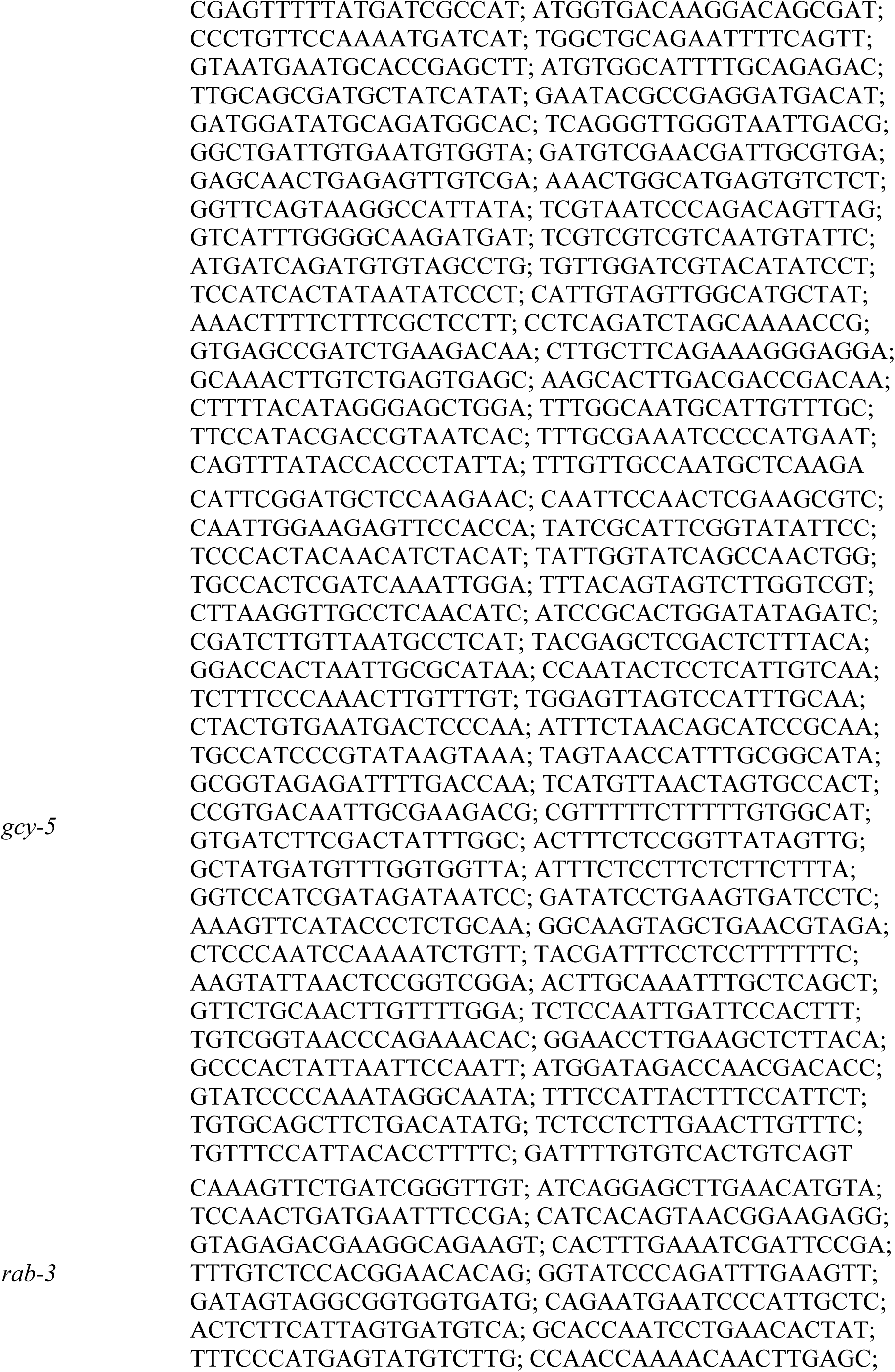
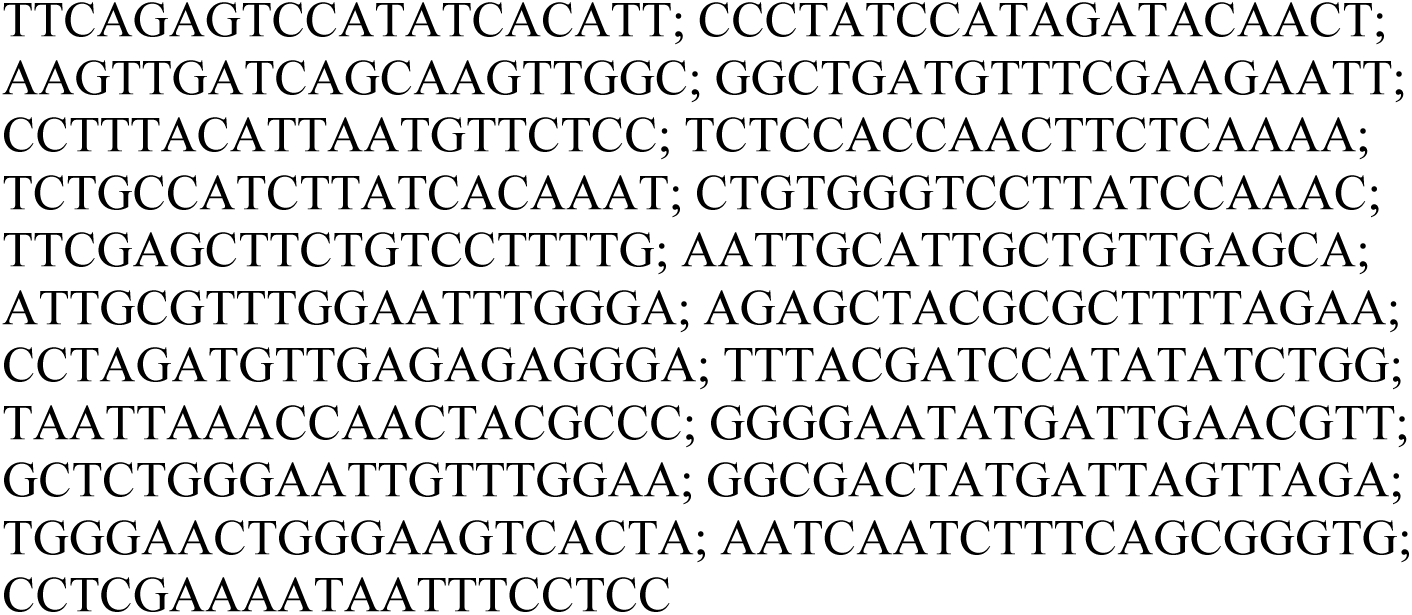
smFISH probes used in this study:

## REFERENCES

Becker, J. S., D. Nicetto and K. S. Zaret (2016). “H3K9me3-Dependent Heterochromatin: Barrier to Cell Fate Changes.” Trends Genet 32(1): 29–41.

Bishop, T., K. W. Lau, A. C. Epstein, S. K. Kim, M. Jiang, D. O’Rourke, C. W. Pugh, J. M. Gleadle, M. S. Taylor, J. Hodgkin and P. J. Ratcliffe (2004). “Genetic analysis of pathways regulated by the von Hippel-Lindau tumor suppressor in Caenorhabditis elegans.” PLoS Biol 2(10): e289.

Brumbaugh, J., B. Di Stefano and K. Hochedlinger (2019). “Reprogramming: identifying the mechanisms that safeguard cell identity.” Development 146(23).

Carey, B. W., L. W. Finley, J. R. Cross, C. D. Allis and C. B. Thompson (2015). “Intracellular alpha-ketoglutarate maintains the pluripotency of embryonic stem cells.” Nature 518(7539): 413–416.

Chang, S., R. J. Johnston, Jr. and O. Hobert (2003). “A transcriptional regulatory cascade that controls left/right asymmetry in chemosensory neurons of C. elegans.” Genes Dev 17(17): 2123–2137.

Cheloufi, S., U. Elling, B. Hopfgartner, Y. L. Jung, J. Murn, M. Ninova, M. Hubmann, A. I. Badeaux, C. Euong Ang, D. Tenen, D. J. Wesche, N. Abazova, M. Hogue, N. Tasdemir, J. Brumbaugh, P. Rathert, J. Jude, F. Ferrari, A. Blanco, M. Fellner, D. Wenzel, M. Zinner, S. E. Vidal, O. Bell, M. Stadtfeld, H. Y. Chang, G. Almouzni, S. W. Lowe, J. Rinn, M. Wernig, A. Aravin, Y. Shi, P. J. Park, J. M. Penninger, J. Zuber and K. Hochedlinger (2015). “The histone chaperone CAF-1 safeguards somatic cell identity.” Nature 528(7581): 218–224.

Chen, X. and J. Ding (2023). “Molecular insights into the catalysis and regulation of mammalian NAD-dependent isocitrate dehydrogenases.” Curr Opin Struct Biol 82: 102672.

Chin, R. M., X. Fu, M. Y. Pai, L. Vergnes, H. Hwang, G. Deng, S. Diep, B. Lomenick, V. S. Meli, G. C. Monsalve, E. Hu, S. A. Whelan, J. X. Wang, G. Jung, G. M. Solis, F. Fazlollahi, C. Kaweeteerawat, A. Quach, M. Nili, A. S. Krall, H. A. Godwin, H. R. Chang, K. F. Faull, F. Guo, M. Jiang, S. A. Trauger, A. Saghatelian, D. Braas, H. R. Christofk, C. F. Clarke, M. A. Teitell, M. Petrascheck, K. Reue, M. E. Jung, A. R. Frand and J. Huang (2014). “The metabolite alpha-ketoglutarate extends lifespan by inhibiting ATP synthase and TOR.” Nature 510(7505): 397–401.

Davis, R. L., H. Weintraub and A. B. Lassar (1987). “Expression of a single transfected cDNA converts fibroblasts to myoblasts.” Cell 51(6): 987–1000.

DeBerardinis, R. J., A. Mancuso, E. Daikhin, I. Nissim, M. Yudkoff, S. Wehrli and C. B. Thompson (2007). “Beyond aerobic glycolysis: transformed cells can engage in glutamine metabolism that exceeds the requirement for protein and nucleotide synthesis.” Proc Natl Acad Sci U S A 104(49): 19345–19350.

Epstein, A. C., J. M. Gleadle, L. A. McNeill, K. S. Hewitson, J. O’Rourke, D. R. Mole, M. Mukherji, E. Metzen, M. I. Wilson, A. Dhanda, Y. M. Tian, N. Masson, D. L. Hamilton, P. Jaakkola, R. Barstead, J. Hodgkin, P. H. Maxwell, C. W. Pugh, C. J. Schofield and P. J. Ratcliffe (2001). “C. elegans EGL-9 and mammalian homologs define a family of dioxygenases that regulate HIF by prolyl hydroxylation.” Cell 107(1): 43–54.

Etchberger, J. F., A. Lorch, M. C. Sleumer, R. Zapf, S. J. Jones, M. A. Marra, R. A. Holt, D. G. Moerman and O. Hobert (2007). “The molecular signature and cis-regulatory architecture of a C. elegans gustatory neuron.” Genes Dev 21(13): 1653–1674.

Feng, T., A. Yamamoto, S. E. Wilkins, E. Sokolova, L. A. Yates, M. Munzel, P. Singh, R. J. Hopkinson, R. Fischer, M. E. Cockman, J. Shelley, D. C. Trudgian, J. Schodel, J. S. McCullagh, W. Ge, B. M. Kessler, R. J. Gilbert, L. Y. Frolova, E. Alkalaeva, P. J. Ratcliffe, C. J. Schofield and M. L. Coleman (2014). “Optimal translational termination requires C4 lysyl hydroxylation of eRF1.” Mol Cell 53(4): 645–654.

Figueroa, M. E., O. Abdel-Wahab, C. Lu, P. S. Ward, J. Patel, A. Shih, Y. Li, N. Bhagwat, A. Vasanthakumar, H. F. Fernandez, M. S. Tallman, Z. Sun, K. Wolniak, J. K. Peeters, W. Liu, S. E. Choe, V. R. Fantin, E. Paietta, B. Lowenberg, J. D. Licht, L. A. Godley, R. Delwel, P. J. Valk, C. B. Thompson, R. L. Levine and A. Melnick (2010). “Leukemic IDH1 and IDH2 mutations result in a hypermethylation phenotype, disrupt TET2 function, and impair hematopoietic differentiation.” Cancer Cell 18(6): 553–567.

Folmes, C. D., T. J. Nelson, A. Martinez-Fernandez, D. K. Arrell, J. Z. Lindor, P. P. Dzeja, Y. Ikeda, C. Perez-Terzic and A. Terzic (2011). “Somatic oxidative bioenergetics transitions into pluripotency-dependent glycolysis to facilitate nuclear reprogramming.” Cell Metab 14(2): 264–271.

Gabriel, J. L., P. R. Zervos and G. W. Plaut (1986). “Activity of purified NAD-specific isocitrate dehydrogenase at modulator and substrate concentrations approximating conditions in mitochondria.” Metabolism 35(7): 661–667.

Hajduskova, M., G. Baytek, E. Kolundzic, A. Gosdschan, M. Kazmierczak, A. Ofenbauer, M. L. Beato Del Rosal, S. Herzog, N. Ul Fatima, P. Mertins, S. Seelk-Muthel and B. Tursun (2019). “MRG-1/MRG15 Is a Barrier for Germ Cell to Neuron Reprogramming in Caenorhabditis elegans.” Genetics 211(1): 121– 139.

Jin, Y., R. Hoskins and H. R. Horvitz (1994). “Control of type-D GABAergic neuron differentiation by C. elegans UNC-30 homeodomain protein.” Nature 372(6508): 780–783.

Kaelin, W. G., Jr. and P. J. Ratcliffe (2008). “Oxygen sensing by metazoans: the central role of the HIF hydroxylase pathway.” Mol Cell 30(4): 393–402.

Kazmierczak, M., I. D. C. Farre, A. Ofenbauer, S. Herzog and B. Tursun (2021). “The CONJUDOR pipeline for multiplexed knockdown of gene pairs identifies RBBP-5 as a germ cell reprogramming barrier in C. elegans.” Nucleic Acids Res 49(4): e22.

Kim, W., R. S. Underwood, I. Greenwald and D. D. Shaye (2018). “OrthoList 2: A New Comparative Genomic Analysis of Human and Caenorhabditis elegans Genes.” Genetics 210(2): 445–461.

Kolundzic, E., A. Ofenbauer, S. I. Bulut, B. Uyar, G. Baytek, A. Sommermeier, S. Seelk, M. He, A. Hirsekorn, D. Vucicevic, A. Akalin, S. Diecke, S. A. Lacadie and B. Tursun (2018). “FACT Sets a Barrier for Cell Fate Reprogramming in Caenorhabditis elegans and Human Cells.” Dev Cell 46(5): 611–626 e612.

Krieg, A. J., E. B. Rankin, D. Chan, O. Razorenova, S. Fernandez and A. J. Giaccia (2010). “Regulation of the histone demethylase JMJD1A by hypoxia-inducible factor 1 alpha enhances hypoxic gene expression and tumor growth.” Mol Cell Biol 30(1): 344–353.

Lin, Z.-H., S.-Y. Chang, W.-C. Shen, Y.-H. Lin, C.-L. Shen, S.-B. Liao, Y.-C. Liu, C.-S. Chen, T.-T. Ching and H.-D. Wang (2022). “Isocitrate Dehydrogenase Alpha-1 Modulates Lifespan and Oxidative Stress Tolerance in Caenorhabditis elegans.” International Journal of Molecular Sciences 24(1).

Lu, C., P. S. Ward, G. S. Kapoor, D. Rohle, S. Turcan, O. Abdel-Wahab, C. R. Edwards, R. Khanin, M. E. Figueroa, A. Melnick, K. E. Wellen, D. M. O’Rourke, S. L. Berger, T. A. Chan, R. L. Levine, I. K. Mellinghoff and C. B. Thompson (2012). “IDH mutation impairs histone demethylation and results in a block to cell differentiation.” Nature 483(7390): 474–478.

Onder, T. T., N. Kara, A. Cherry, A. U. Sinha, N. Zhu, K. M. Bernt, P. Cahan, B. O. Marcarci, J. Unternaehrer, P. B. Gupta, E. S. Lander, S. A. Armstrong and G. Q. Daley (2012). “Chromatin-modifying enzymes as modulators of reprogramming.” Nature 483(7391): 598–602.

Owen, O. E., S. C. Kalhan and R. W. Hanson (2002). “The key role of anaplerosis and cataplerosis for citric acid cycle function.” J Biol Chem 277(34): 30409–30412.

Patel, T., B. Tursun, D. P. Rahe and O. Hobert (2012). “Removal of Polycomb repressive complex 2 makes C. elegans germ cells susceptible to direct conversion into specific somatic cell types.” Cell Rep 2(5): 1178–1186.

Pollard, P. J., C. Loenarz, D. R. Mole, M. A. McDonough, J. M. Gleadle, C. J. Schofield and P. J. Ratcliffe (2008). “Regulation of Jumonji-domain-containing histone demethylases by hypoxia-inducible factor (HIF)-1alpha.” Biochem J 416(3): 387–394.

Schneuwly, S., R. Klemenz and W. J. Gehring (1987). “Redesigning the Body Plan of Drosophila by Ectopic Expression of the Homeotic Gene Antennapedia.” Nature 325(6107): 816–818.

Shen, C., D. Nettleton, M. Jiang, S. K. Kim and J. A. Powell-Coffman (2005). “Roles of the HIF-1 hypoxia-inducible factor during hypoxia response in Caenorhabditis elegans.” J Biol Chem 280(21): 20580–20588.

Soufi, A. and K. S. Zaret (2013). “Understanding impediments to cellular conversion to pluripotency by assessing the earliest events in ectopic transcription factor binding to the genome.” Cell Cycle 12(10): 1487–1491.

Takahashi, K. and S. Yamanaka (2006). “Induction of pluripotent stem cells from mouse embryonic and adult fibroblast cultures by defined factors.” Cell 126(4): 663–676.

TeSlaa, T., A. C. Chaikovsky, I. Lipchina, S. L. Escobar, K. Hochedlinger, J. Huang, T. G. Graeber, D. Braas and M. A. Teitell (2016). “alpha-Ketoglutarate Accelerates the Initial Differentiation of Primed Human Pluripotent Stem Cells.” Cell Metab 24(3): 485–493.

Tischler, J., W. H. Gruhn, J. Reid, E. Allgeyer, F. Buettner, C. Marr, F. Theis, B. D. Simons, L. Wernisch and M. A. Surani (2019). “Metabolic regulation of pluripotency and germ cell fate through alpha-ketoglutarate.” EMBO J 38(1).

Tursun, B., T. Patel, P. Kratsios and O. Hobert (2011). “Direct conversion of C. elegans germ cells into specific neuron types.” Science 331(6015): 304–308.

Vierbuchen, T., A. Ostermeier, Z. P. Pang, Y. Kokubu, T. C. Sudhof and M. Wernig (2010). “Direct conversion of fibroblasts to functional neurons by defined factors.” Nature 463(7284): 1035–1041.

Wellmann, S., M. Bettkober, A. Zelmer, K. Seeger, M. Faigle, H. K. Eltzschig and C. Buhrer (2008). “Hypoxia upregulates the histone demethylase JMJD1A via HIF-1.” Biochem Biophys Res Commun 372(4): 892–897.

Xiao, D., L. Zeng, K. Yao, X. Kong, G. Wu and Y. Yin (2016). “The glutamine-alpha-ketoglutarate (AKG) metabolism and its nutritional implications.” Amino Acids 48(9): 2067–2080.

Xiao, M., H. Yang, W. Xu, S. Ma, H. Lin, H. Zhu, L. Liu, Y. Liu, C. Yang, Y. Xu, S. Zhao, D. Ye, Y. Xiong and K. L. Guan (2012). “Inhibition of alpha-KG-dependent histone and DNA demethylases by fumarate and succinate that are accumulated in mutations of FH and SDH tumor suppressors.” Genes Dev 26(12): 1326–1338.

Xu, W., H. Yang, Y. Liu, Y. Yang, P. Wang, S. H. Kim, S. Ito, C. Yang, P. Wang, M. T. Xiao, L. X. Liu, W. Q. Jiang, J. Liu, J. Y. Zhang, B. Wang, S. Frye, Y. Zhang, Y. H. Xu, Q. Y. Lei, K. L. Guan, S. M. Zhao and Y. Xiong (2011). “Oncometabolite 2-hydroxyglutarate is a competitive inhibitor of alpha-ketoglutarate-dependent dioxygenases.” Cancer Cell 19(1): 17–30.

Yang, C., B. Ko, C. T. Hensley, L. Jiang, A. T. Wasti, J. Kim, J. Sudderth, M. A. Calvaruso, L. Lumata, M. Mitsche, J. Rutter, M. E. Merritt and R. J. DeBerardinis (2014). “Glutamine oxidation maintains the TCA cycle and cell survival during impaired mitochondrial pyruvate transport.” Mol Cell 56(3): 414–424.

Zhang, D., Y. Wang, Z. Shi, J. Liu, P. Sun, X. Hou, J. Zhang, S. Zhao, B. P. Zhou and J. Mi (2015). “Metabolic reprogramming of cancer-associated fibroblasts by IDH3alpha downregulation.” Cell Rep 10(8): 1335–1348.

Zhang, J., E. Nuebel, G. Q. Daley, C. M. Koehler and M. A. Teitell (2012). “Metabolic regulation in pluripotent stem cells during reprogramming and self-renewal.” Cell Stem Cell 11(5): 589–595.

Zhuang, Q., T. Feng and M. L. Coleman (2015). “Modifying the maker: Oxygenases target ribosome biology.” Translation (Austin) 3(1): e1009331.

## References

Antoniewicz, M. R. (2018). A guide to ^13C metabolic flux analysis for the cancer biologist. Experimental & Molecular Medicine, 50(4), 1–13. doi:10.1038/s12276-018-0060-y.

Bettinger, J. C., Lee, K., & Rougvie, A. E. (1996). Stage-specific accumulation of the terminal differentiation factor LIN-29 during *Caenorhabditis elegans* development. Development, 122(8), 2517–2527. doi:10.1242/dev.122.8.2517.

Brenner, S. (1974). The genetics of *Caenorhabditis elegans*. Genetics, 77(1), 71–94. doi:10.1093/genetics/77.1.71.

Chin, R. M., Fu, X., Pai, M. Y., Vergnes, L., Hwang, H., Deng, G., Diep, S., Lomenick, B., Meli, V. S., Monsalve, G. C., Hu, E., Whelan, S. A., Wang, J. X., Jung, G., Solis, G. M., Fazlollahi, F., Kaweeteerawat, C., Quach, A., Nili, M., … Huang, J. (2014). The metabolite α-ketoglutarate extends lifespan by inhibiting ATP synthase and TOR. Nature, 510(7505), 397–401. doi:10.1038/nature13264.

Dokshin, G. A., Ghanta, K. S., Piscopo, K. M., & Mello, C. C. (2018). Robust genome editing with short single-stranded and long, partially single-stranded DNA donors in *Caenorhabditis elegans*. Genetics, 210(3), 781–787. doi:10.1534/genetics.118.301532.

Jones, A. R., Francis, R., & Schedl, T. (1996). GLD-1, a cytoplasmic protein essential for oocyte differentiation, shows stage- and sex-specific expression during *Caenorhabditis elegans* germline development. Developmental Biology, 180(1), 165–183. doi:10.1006/dbio.1996.0293.

Kamath, R. S., Fraser, A. G., Dong, Y., Poulin, G., Durbin, R., Gotta, M., Kanapin, A., Le Bot, N., Moreno, S., Sohrmann, M., Welchman, D. P., Zipperlen, P., & Ahringer, J. (2003). Systematic functional analysis of the *Caenorhabditis elegans* genome using RNAi. Nature, 421(6920), 231–237. doi:10.1038/nature01278.

Kempa, S., Hummel, J., Schwemmer, T., Pietzke, M., Strehmel, N., Wienkoop, S., Kopka, J., & Weckwerth, W. (2009). An automated GCxGC-TOF-MS protocol for batch-wise extraction and alignment of mass isotopomer matrixes from differential ^13C-labelling experiments: A case study for photoautotrophic-mixotrophic grown *Chlamydomonas reinhardtii* cells. Journal of Basic Microbiology, 49(1), 82–91. doi:10.1002/jobm.200800337.

Kolundzic, E., Ofenbauer, A., Bulut, S. I., Uyar, B., Baytek, G., Sommermeier, A., Seelk, S., He, M., Hirsekorn, A., Vucicevic, D., Akalin, A., Diecke, S., Lacadie, S. A., & Tursun, B. (2018). FACT sets a barrier for cell fate reprogramming in *Caenorhabditis elegans* and human cells. Developmental Cell, 46(5), 611–626.e12. doi:10.1016/j.devcel.2018.07.006.

Kuich, P. H. J. L., Hoffmann, N., & Kempa, S. (2015). Maui-VIA: A user-friendly software for visual identification, alignment, correction, and quantification of gas chromatography– mass spectrometry data. Frontiers in Bioengineering and Biotechnology, 2, 84. doi:10.3389/fbioe.2014.00084.

Pietzke, M., Zasada, C., Mudrich, S., & Kempa, S. (2014). Decoding the dynamics of cellular metabolism and the action of 3-bromopyruvate and 2-deoxyglucose using pulsed stable isotope-resolved metabolomics. Cancer & Metabolism, 2, 9. doi:10.1186/2049-3002-2-9.

Roessner-Tunali, U., Hegemann, B., Lytovchenko, A., Carrari, F., Bruedigam, C., Granot, D., & Fernie, A. R. (2003). Metabolic profiling of transgenic tomato plants overexpressing hexokinase reveals that the influence of hexose phosphorylation diminishes during fruit development. Plant Physiology, 133(1), 84–99. doi:10.1104/pp.103.023572.

Sarasija, S., & Norman, K. R. (2015). A γ-secretase independent role for presenilin in calcium homeostasis impacts mitochondrial function and morphology in *Caenorhabditis elegans*. Genetics, 201(4), 1453–1466. doi:10.1534/genetics.115.182808.

Sarasija, S., & Norman, K. R. (2018). Analysis of mitochondrial structure in the body wall muscle of *Caenorhabditis elegans*. Bio-protocol, 8(7), e2801. doi:10.21769/BioProtoc.2801.

Tursun, B., Patel, T., Kratsios, P., & Hobert, O. (2011). Direct conversion of *C. elegans* germ cells into specific neuron types. Science, 331(6015), 304–308. doi:10.1126/science.1199082.

